# Cooperativity and additivity in plant enhancers

**DOI:** 10.1101/2023.11.21.568158

**Authors:** Tobias Jores, Jackson Tonnies, Nicholas A. Mueth, Andrés Romanowski, Stanley Fields, Josh T. Cuperus, Christine Queitsch

**Author notes:** Authors for correspondence (T.J.), (C.Q.). These authors contributed equally (Tobias Jores, Jackson Tonnies). The authors responsible for distribution of materials integral to the findings presented in this article in accordance with the policy described in the Instructions for Authors (https://academic.oup.com/plcell/pages/General-Instructions) are: Tobias Jores and Christine Queitsch.

## Abstract

Enhancers are *cis*-regulatory elements that shape gene expression in response to numerous developmental and environmental cues. In animals, several models have been proposed to explain how enhancers integrate the activity of multiple transcription factors. However, it remains largely unknown how plant enhancers integrate transcription factor activity. Here, we use Plant STARR-seq to characterize three light-responsive plant enhancers—*AB80*, *Cab-1*, and *rbcS-E9*—derived from genes active in photosynthesis. Saturation mutagenesis reveals mutations, many of which cluster in short regions, that strongly reduce enhancer activity in the light, in the dark or in both conditions. When tested in the light, these mutation-sensitive regions do not function on their own; rather, cooperative interactions with other such regions are required for full activity. Epistatic interactions occur between mutations in adjacent mutation-sensitive regions, and the spacing and order of mutation-sensitive regions in synthetic enhancers affects enhancer activity. In contrast, when tested in the dark, mutation-sensitive regions act independently and additively in conferring enhancer activity. Taken together, this work demonstrates that plant enhancers show evidence for both cooperative and additive interactions among their functional elements. This knowledge can be harnessed to design strong, condition-specific synthetic enhancers.

## Introduction

Enhancers play a pivotal role in orchestrating the precise gene expression programs required for plants to develop and thrive in concert with their ever-changing abiotic and biotic environments (Schmitz et al., 2022; Marand et al., 2023). They integrate spatiotemporal, developmental, and environmental cues by binding to transcription factors to enhance the transcription of their target genes.

Three models have been proposed to explain how enhancers in animal cells integrate the activity of multiple transcription factors (Arnosti and Kulkarni, 2005; Spitz and Furlong, 2012; Jindal and Farley, 2021; Kim and Wysocka, 2023). In the “billboard” model, the binding or activity of individual transcription factors is not dependent on other factors binding to the same enhancer (Kulkarni and Arnosti, 2003; de Boer et al., 2020). In contrast, in the “enhanceosome” model, enhancers recruit a highly ordered transcription factor complex (Thanos and Maniatis, 1995; Panne, 2008). Full activity is reached only in the presence of all complex members, and enhancer activity is drastically reduced if even a single transcription factor is missing or if the enhancer grammar—the orientation, spacing, and order of the individual transcription factor binding sites—is altered. Finally, the “transcription factor collective” model describes enhancers that recruit a complex composed of multiple transcription factors—all of which are necessary for full activity—without requiring a fixed enhancer grammar (Junion et al., 2012; Uhl et al., 2016). The three models differ in two main aspects: their requirement for a specific enhancer grammar (high for the enhanceosome model; low for the billboard and transcription factor collective models) and the degree of cooperativity between the recruited transcription factors (high for the enhanceosome and transcription factor collective models; low for the billboard model). Hence, grammar and cooperativity are key characteristics of enhancers (Kim et al., 2022; Friedman et al., 2023; Song et al., 2023).

To date, only a few enhancers in plants have been well characterized (Weber et al., 2016; Schmitz et al., 2022). Like their animal counterparts, plant enhancers function independently of orientation, are active over a wide range of distances, and occur upstream or downstream of their target promoter (Weber et al., 2016; Schmitz et al., 2022; Marand et al., 2023). Despite these similarities, enhancers in plants and animals differ in their histone modifications (Lu et al., 2019; Yan et al., 2019; Silver et al., 2023). The absence in plants of the canonical histone modifications that mark active enhancers in animals is consistent with the absence in plants of the bi-directional, short-lived enhancer RNAs found in animals (Erhard et al., 2015; Hetzel et al., 2016; Thieffry et al., 2020; Mcdonald et al., 2023; Silver et al., 2023). Thus, plants appear to use different mechanisms than animals for maintaining enhancer activity.

Promoter-bashing approaches and testing of synthetic promoters assembled from transcription factor binding sites have generated evidence for cooperative interactions between functional sub-elements in plants (Benfey et al., 1990; Walcher and Nemhauser, 2012; Cai et al., 2020; Wang et al., 2021). However, we sought to interrogate enhancers at nucleotide resolution with saturation mutagenesis. We therefore took advantage of the Plant STARR-seq method (Jores et al., 2020, 2023) to characterize the functional underpinnings of three light-responsive plant enhancers: *AB80* from the pea *Pisum sativum*, *Cab-1* from the wheat *Triticum aestivum*, and *rbcS-E9* from *P. sativum*. We identify several mutation-sensitive regions within each enhancer that are crucial for condition-specific activity. In the light, strong cooperativity between these regions is displayed, whereas in the dark the mutation-sensitive regions that contribute to enhancer activity function largely independent of each other.

## Results

### Enhancers from photosynthesis genes show light-responsive activity

We used Plant STARR-seq to dissect three enhancers that are associated with photosynthesis genes and possess light-responsive activity (Fluhr et al., 1986; Simpson et al., 1986; Nagy et al., 1987). The *AB80* and *Cab-1* enhancers drive the expression of chlorophyl a-b binding proteins, and the *rbcS-E9* enhancer regulates the expression of a small subunit of the ribulose-1,5-bisphosphate carboxylase. We previously demonstrated that Plant STARR-seq detects the light-responsive activity of these three enhancers (Jores et al., 2020).

We sought to carry out complete saturation mutagenesis of the three enhancers using array synthesis, which limits the length of oligonucleotides that can be synthesized. Therefore, we first tested whether 169-bp long segments, that are amenable to array synthesis, derived from the 5′ or 3′ end of each enhancer show light-responsive activity. We cloned the full-length (as defined by restriction enzyme-based truncation analysis in: Fluhr et al., 1986; Simpson et al., 1986; Nagy et al., 1987) and truncated enhancer segments upstream of the 35S minimal promoter controlling the expression of a barcoded green fluorescent protein (GFP) reporter gene and subjected the pooled library to Plant STARR-seq in tobacco (*Nicotiana benthamiana*) leaves (Figure 1A). To test for light-responsive enhancer activity, we kept the transiently transformed tobacco plants in either normal light/dark cycles (16 h light and 8 h dark) or completely in the dark for two days prior to RNA extraction. While the 35S enhancer, included as a control, showed strong and largely light-independent activity, the activity of the three plant enhancers was light-dependent (Figure 1, B and C).

**Figure 1.**
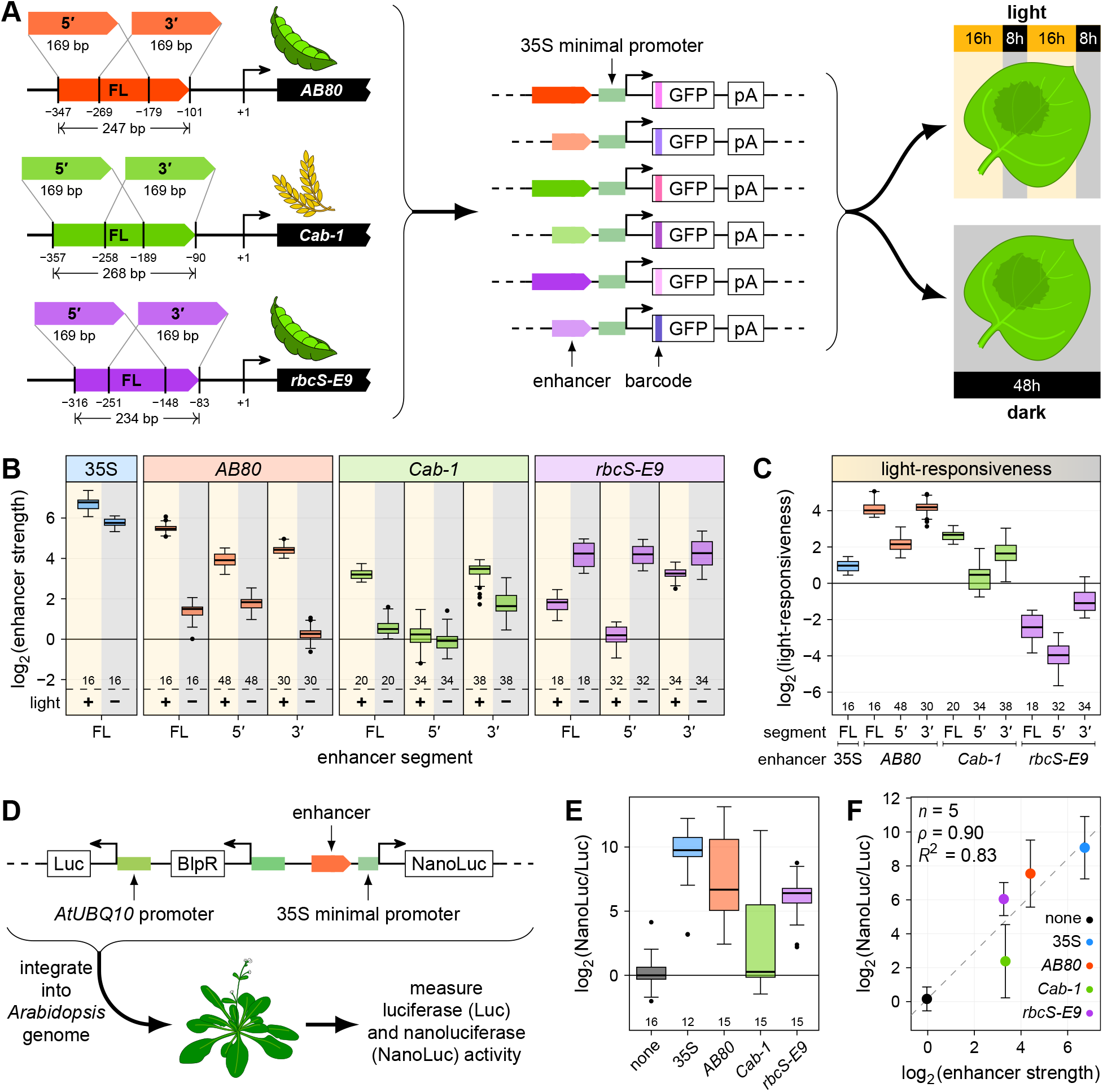
Enhancers from photosynthesis genes show light-responsive activity. **A** and **B**, Full-length (FL) enhancers, as well as 169-bp long segments from their 5′ or 3′ end, of the *Pisum sativum AB80* and *rbcS-E9* genes and the *Triticum aestivum Cab-1* gene were cloned upstream of the 35S minimal promoter driving the expression of a barcoded GFP reporter gene (**A**). All constructs were pooled and the viral 35S enhancer was added as an internal control. The pooled enhancer library was subjected to Plant STARR-seq in tobacco leaves with plants grown for 2 days in normal light/dark cycles (+ light) or completely in the dark (− light) prior to RNA extraction (**B**). Enhancer strength was normalized to a control construct without an enhancer (log_2_ set to 0). **C**, Light-responsiveness (log_2_[enhancer strength^light^/enhancer strength^dark^]) was determined for the indicated enhancer segments. **D**, Transgenic Arabidopsis lines were generated with T-DNAs harboring a constitutively expressed luciferase (Luc) gene and a nanoluciferase (NanoLuc) gene under control of a 35S minimal promoter coupled to the 35S enhancer or the 3′ segments of the *AB80*, *Cab-1*, or *rbcS-E9* enhancers. **E**, Nanoluciferase activity was measured in 5 T2 plants from these lines and normalized to the activity of luciferase. The NanoLuc/Luc ratio was normalized to a control construct without an enhancer (none; log_2_ set to 0). **F**, The mean NanoLuc/Luc ratio was compared to the mean enhancer strength determined by STARR-seq. Pearson’s *R*^2^, Spearman’s ρ, and number (*n*) of enhancers are indicated. A linear regression line is shown as a dashed line. Error bars represent the 95% confidence interval. Box plots in **B**, **C**, and **E** represent the median (center line), upper and lower quartiles (box limits), 1.5× interquartile range (whiskers), and outliers (points) for all corresponding barcodes (**B** and **C**) or plant lines (**E**) from two (**B** and **C**) or three (**E**) independent replicates. Numbers at the bottom of each box plot indicate the number of samples in each group.

In the light, the 5′ and 3′ segments of the *AB80* enhancer were active, albeit not to the extent of the full-length enhancer. In the dark, the *AB80* 5′ segment retained the weak activity of the full-length enhancer, whereas its 3′ segment was inactive (Figure 1B). In the light, the *Cab-1* 3′ segment showed similar strength compared to the full-length enhancer; however, in the dark this segment showed 2-fold higher activity than the full-length enhancer did. The *Cab-1* 5′ segment showed no enhancer activity in either condition (Figure 1B). The *rbcS-E9* enhancer showed stronger activity in the dark than in the light, and this activity was retained in both the 5′ and 3′ segments. However, in the light, the *rbcS-E9* 3′ segment showed approximately 3-fold more activity than the full-length enhancer, whereas the 5′ segment was inactive (Figure 1B).

While the 169-bp enhancer segments were generally weaker than their full-length counterparts, this was not always the case. The *Cab-1* 3’ segment showed higher activity than the full-length enhancer in the dark and the *rbcS-E9* 3’ segment showed higher activity than the full-length enhancer in the light (Figure 1B). These observations suggest that the *Cab-1* and *rbcS-E9* enhancers also harbor putative repressive elements. These repressive elements reside outside of the 3’ enhancer segments in both enhancers.

Enhancer constructs were tested both in the forward and the reverse orientation because enhancers are expected to function in an orientation-independent manner (Banerji et al., 1981; Fang et al., 1989; Jores et al., 2020; Schmitz et al., 2022). Indeed, we observed this property for the full-length and truncated plant enhancers assayed here (Supplemental Figure S1). At least two biological replicates were performed for all experiments, and the results were highly reproducible (*R*^2^ ≥ 0.87; Supplemental Figure S2).

Finally, we confirmed that the 169-bp enhancer segments active in the transient Plant STARR-seq assay also show enhancer activity when assayed in stable transgenic lines. We cloned the 3′ segments of the *AB80*, *Cab-1*, and *rbcS-E9* enhancers upstream of a 35S minimal promoter driving the expression of nanoluciferase. The T-DNA also contained a luciferase gene controlled by the constitutive *Arabidopsis thaliana POLYUBIQUITIN 10* promoter to control for effects on expression due to genomic context (Figure 1D). We generated transgenic *Arabidopsis* Col-0 lines harboring these T-DNAs and measured nanoluciferase and luciferase activities in T2 plants from five independent lines. We detected high enhancer activity (indicated by a high nanoluciferase/luciferase ratio) for the 3′ segments of the *AB80* and *rbcS-E9* enhancers in *Arabidopsis*. However, the 3′ segment of the *Cab-1* enhancer showed low activity (Figure 1E). Shifting the transgenic *Arabidopsis* plants to the dark for four days prior to sample collection did not change the observed nanoluciferase/luciferase ratios (Supplemental Figure S3), presumably because nanoluciferase protein turnover is low *in planta*. Nonetheless, we observed a strong correlation between the Plant STARR-seq and dual-luciferase assays in the light (Figure 1F).

In summary, except for the 5′ segment of *Cab-1*, the 169-bp enhancer segments function as enhancers in at least one light regime, either upregulated in the light or upregulated in the dark (Figure 1B). Therefore, the shortened enhancer segments can be used to study the condition-specific activity of plant enhancers.

### Plant enhancers contain multiple mutation-sensitive regions

To identify regions of the *AB80*, *Cab-1*, and *rbcS-E9* enhancers that confer enhancer activity, we subjected the shortened enhancer segments to saturation mutagenesis. We array-synthesized all possible single-nucleotide substitution, deletion, and insertion variants of the 5′ and 3′ segments of the three enhancers, and measured their enhancer strength with Plant STARR-seq in the light and the dark. We determined the activity of over 99% of all possible single-nucleotide variants (Supplemental Data Set 1). While most mutations had little to no effect, some of the variants resulted in an up to four-fold increase or to a decrease to as low as 1/20 in enhancer strength relative to the wild-type sequence (Supplemental Figure S4).

Mutations in all three enhancers with a negative effect on enhancer strength often clustered together, revealing mutation-sensitive regions (Figure 2; Supplemental Figure S5). The effect sizes of mutations within the mutation-sensitive regions in each enhancer were consistent with our observations on the light-responsive activity of the respective wild-type enhancer segments (Figure 1B). For the *AB80* and *Cab-1* enhancers, the effect sizes of mutations within the mutation-sensitive regions were much greater in the light than in the dark (Figure 2, A and B). For the *rbcS-E9* enhancer, mutation-sensitive regions in the dark overlapped between the 5’ and 3’ segments, while mutation-sensitive regions in the light resided both in the overlap region and closer to the 3’ end (Figure 2C). These observations are consistent with our findings that both segments showed similar enhancer strength in the dark compared to the full-length enhancer, whereas only the 3’ segment was active in the light (Figure 1B).

**Figure 2.**
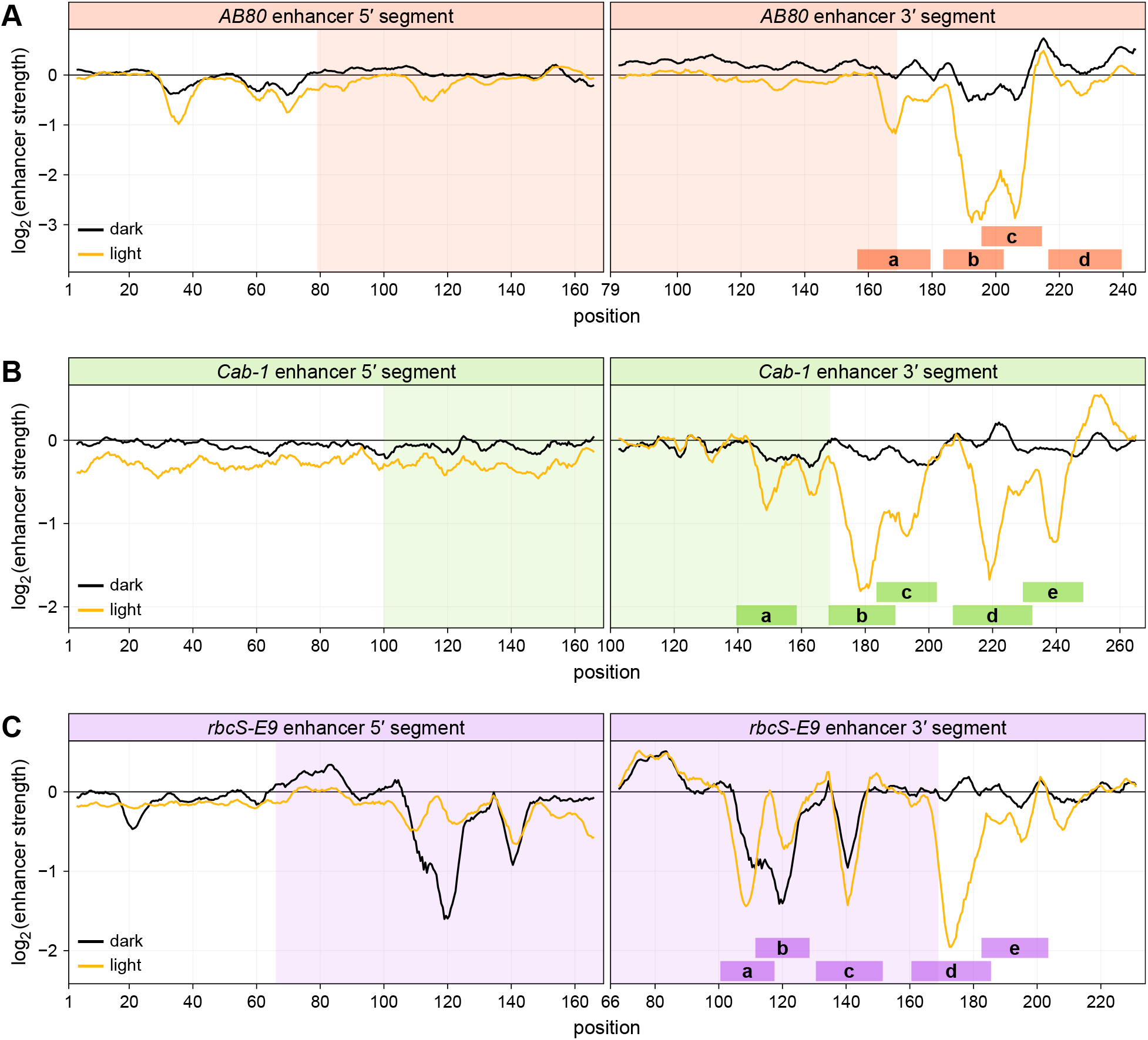
The *AB80*, *Cab-1*, and *rbcS-E9* enhancers contain multiple mutation-sensitive regions. **A**–**C**, All possible single-nucleotide substitution, deletion, and insertion variants of the 5′ and 3′ segments of the *AB80* (**A**), *Cab-1* (**B**), and *rbcS-E9* (**C**) enhancers were subjected to Plant STARR-seq in tobacco plants grown in normal light/dark cycles (light) or completely in the dark (dark) for two days prior to RNA extraction. Enhancer strength was normalized to the wild-type variant (log_2_ set to 0). A sliding average (window size = 6 bp) of the mean enhancer strength for all variants at a given position is shown. The shaded area indicates the region where the 5′ and 3′ segments overlap. Mutation-sensitive regions in the 3′ enhancer segments are indicated by shaded rectangles labeled a–e.

The 5′ and 3′ segments of each enhancer overlap partially (overlap length: 91 bp for *AB80*, 70 bp for *Cab-1*, and 104 bp for *rbcS-E9*). We asked whether mutations in the overlap region show similar effects when tested in the context of the 5′ or 3′ segment in the light or dark. In the dark, context did not matter for the effects of mutations. For example, the mutation-sensitive regions in the *rbcS-E9* enhancer reside in similar positions (Figure 2C) and mutational effects in the 5′ and 3′ segments of the *rbcS-E9* enhancer strongly correlated (Supplemental Figure S6). Although the *AB80* and *Cab-1* enhancers showed little activity in the dark, the variant effects in their respective overlap regions showed positive correlation in this condition. In contrast, for all three enhancers, correlations between variant effects in the overlap region were lower in the light (Supplemental Figure S6). In the light, both the *Cab-1* and the *rbcS-E9* enhancer harbored mutation-sensitive regions in the parts of the 3′ segment part that overlap with the 5′ segment. However, mutations in the corresponding locations of the 5′ segment were weaker or showed no effect on enhancer function (Figure 2, B and C). We conclude that the presence of a mutation-sensitive region, and the effect sizes of mutations within it, depends on the surrounding sequence context (*i.e.*, whether it was tested in the 5′ or 3′ segment) in the light, suggesting cooperative interactions among individual enhancer regions in this condition.

### Mutation-sensitive regions harbor transcription factor binding sites

Because enhancers function by recruiting transcription factors, we searched the enhancer sequences for matches to known transcription factor binding motifs (O’Malley et al., 2016; Tian et al., 2020; Jores et al., 2021). This approach identified 41 putative binding sites; however, 32 (78%) of them were located outside of mutation-sensitive regions, and 5 of the 14 mutation-sensitive regions were not predicted to contain any transcription factor binding site (Supplemental Figure S7).

As an alternative approach, we leveraged the saturation mutagenesis data to identify sequence motifs that contribute to enhancer function. To this end, we manually selected regions (referred to by lowercase letters from a to e) in the 3′ segments of the *AB80*, *Cab-1*, and *rbcS-E9* enhancers with strong mutational sensitivity (Figure 2 and Figure 3A). We then used the measured enhancer strength in the light for all possible single-nucleotide substitution variants in these regions to generate sequence logo plots, which show position-specific nucleotide preferences associated with enhancer strength. All but one of the motifs generated in this way matched known transcription factor binding sites (Figure 3, B–D). Putative binding sites for MYB family transcription factors are found in *AB80* regions a and d, in *Cab-1* regions b and c, and in *rbcS-E9* regions b, c, and e. A G-box motif, a potential binding site for bHLH and bZip transcription factors (O’Malley et al., 2016), is found in *AB80* region b and *rbcS-E9* region a. A putative binding site for TCP family transcription factors is found in *Cab-1* region a. Although no matching transcription factor binding motif was found for *AB80* region c and *Cab-1* region d, both regions contain a mutation-sensitive CCAAT sequence which could be a target of Nuclear Factor Y transcription factors (Gnesutta et al., 2019). Similarly, *rbcS-E9* region d contains a TGTGG pentanucleotide which could be a target of the *CONSTANS* transcription factor and related CCT transcription factors, which form hetero-trimers with Nuclear Factor Y transcription factors (Tiwari et al., 2010; Gnesutta et al., 2017).

**Figure 3.**
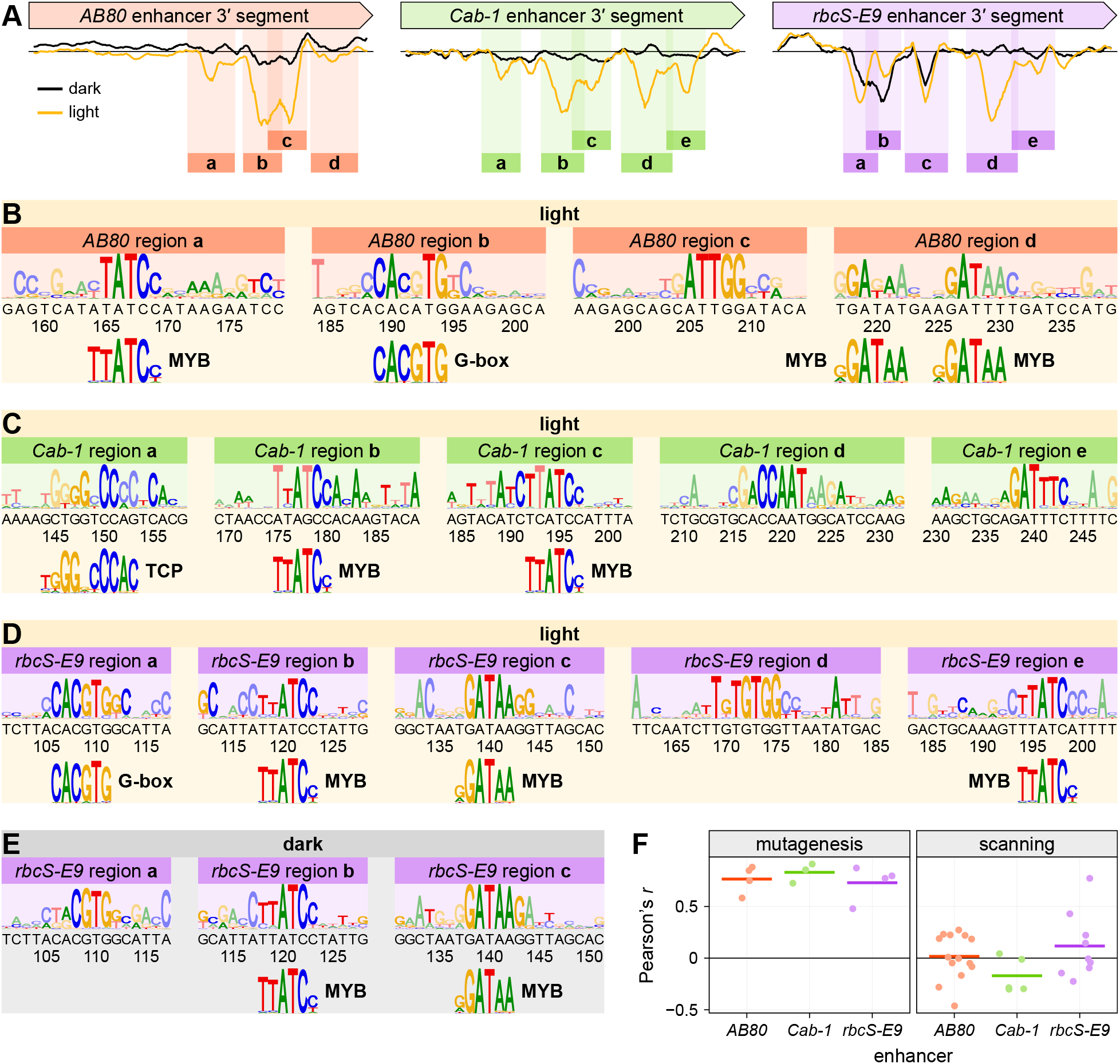
Mutation-sensitive regions harbor transcription factor binding sites. **A**, 4 to 5 mutation-sensitive regions (shaded rectangles; labeled a–e) were defined for the 3′ segments of the *AB80*, *Cab-1*, and *rbcS-E9* enhancers. The mutational sensitivity plots are reproduced from Figure 2. **B**–**E**, Sequence logo plots were generated from the enhancer strength in the light (**B**–**D**) or dark (**E**) of all possible single-nucleotide substitution variants within the indicated mutation-sensitive regions of the *AB80* (**B**), *Cab-1* (**C**), or *rbcS-E9* (**D** and **E**) enhancers. The sequence of the wild-type enhancer and the position along it is shown on the x axis. Letters with dark colors in the logo plot represent wild-type bases. The sequence logos for each region were compared to known transcription factor binding motifs and significant matches are shown below the plots. **F**, For each transcription factor binding motif matching a sequence logo plots derived from the saturation mutagenesis data in the light (mutagenesis; see **B**–**D**) or identified by the motif-scanning approach (scanning; see Supplemental Figure S7), the correlation (Pearson’s *r*) between the strength of an enhancer variant and the score of how well the variant sequence matches this motif is plotted as points. The lines represent the average correlation for all motifs of a given enhancer.

Since the *rbcS-E9* regions a, b, and c also showed mutational sensitivity in the dark, we performed the same analysis for these regions using enhancer strength measurements in the dark (Figure 3E). The ‘dark’ motifs generated for regions b and c are similar to those found in the light. In contrast, for region a, only the last four nucleotides of the G-box were highly sensitive to substitutions in the dark, indicating that this region might be bound by a different transcription factor in the dark than in the light.

Taken together, by leveraging our saturation mutagenesis data, we were able to assign putative transcription factor binding sites to almost all mutation-sensitive regions of the *AB80*, *Cab-1*, and *rbcS-E9* 3′ segments (Figure 3). In contrast, the commonly employed motif-scanning approach did not yield results for several mutation-sensitive regions (Supplemental Figure S7). Moreover, only two (the G-box in *rbcS-E9* region a and the MYB binding site in *rbcS-E9* region b) of the twelve *de novo* motifs based on the saturation mutagenesis data were also identified with the motif-scanning approach.

To test if the motifs identified by the motif-scanning approach are associated with enhancer function, we calculated the correlation between the enhancer strength of a given variant and the score of how well the variant sequence matches the corresponding transcription factor motif. As a positive control, we first determined this correlation for the transcription factor binding motifs inferred from the saturation mutagenesis data. As expected, correlations were high for these motifs. In stark contrast, the motifs from the motif-scanning approach often showed low correlation (Figure 3F). Taken together, these results highlight the power of the saturation mutagenesis-based approach to identify functionally relevant transcription factor binding sites.

We attribute the failure of the motif-scanning approach to the fact that the three enhancer sequences analyzed here often deviate from the consensus transcription factor binding motif by one or two nucleotides. Suboptimal transcription factor binding sites have been observed in animal enhancers, where they ensure precise regulation of enhancer activity by requiring cooperative transcription factors for efficient binding (Farley et al., 2015, 2016). Saturation mutagenesis detects the relevance of such suboptimal motifs by sampling all possible nucleotides.

### The *AB80*, *Cab-1*, and *rbcS-E9* enhancers are regulated by the circadian clock

The *AB80*, *Cab-1*, and *rbcS-E9* enhancers contain putative binding sites for MYB family transcription factors, which have been implicated in the regulation of multiple biological processes, including the circadian clock (Carré and Kim, 2002; Laosuntisuk et al., 2023). Since *Cab-1* is regulated by the circadian clock (Fejes et al., 1990), we asked whether the same is true for the *AB80* and *rbcS-E9* enhancers, and if so, whether we can pinpoint the sequence motifs that mediate this circadian regulation. To address these questions, we used Plant STARR-seq to measure the activity of all single-nucleotide variants of the *AB80*, *Cab-1*, and *rbcS-E9* enhancers in constant light over a time course of 24 hours, with samples taken every 6 hours (Figure 4A). Consistent with the hypothesis that the circadian clock affects the activity of these enhancers, the correlation between Plant STARR-seq samples was highest for samples obtained 24 hours apart from each other and lowest for samples separated by 12 hours (Supplemental Figure S8).

**Figure 4.**
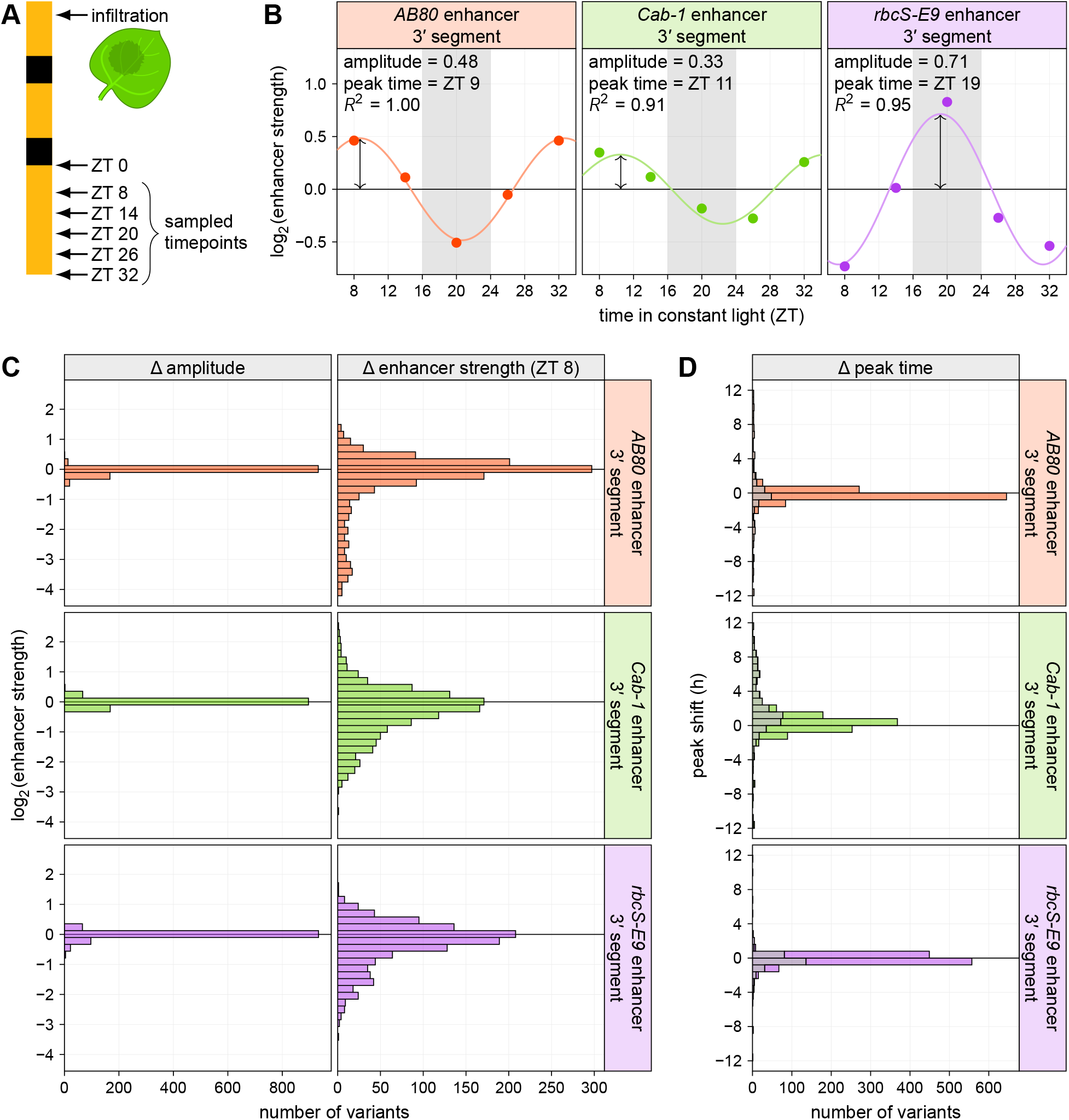
Circadian oscillation is robustly encoded in the *AB80*, *Cab-1*, and *rbcS-E9* enhancers. **A**, All possible single-nucleotide variants of the *AB80*, *Cab-1*, and *rbcS-E9* enhancers were subjected to Plant STARR-seq in tobacco leaves. On the morning of the third day after transformation (ZT 0), the plants were shifted to constant light. Leaves were harvested for RNA extraction starting at mid-day (ZT 8) and in 6 hour intervals (ZT 14, 20, 26, and 32) afterwards. **B**, A sine wave with a period of 24 h was fitted to the enhancer strength of a given variant across all sampled time points. The fitted line is plotted together with the measured data points for the wild-type enhancers. The equilibrium point of the curves was set to 0. The amplitude is shown as a two-sided arrow at the time of highest enhancer strength (peak time). The goodness-of-fit (R^2^) is indicated. The shaded gray area represents the timing of the dark period if the plants had not been shifted to constant light. **C** and **D**, Histograms of the difference between the amplitude (**C**) and peak time (**D**) of each single-nucleotide variant relative to the wild-type enhancer. For comparison, the difference in enhancer strength at ZT 8 is also shown in **C**. Variants with a below average goodness-of-fit are grayed out in **D**. Only data for the 3′ enhancer segments is shown.

The activity of all three enhancers was influenced by the circadian clock. The *AB80* and *Cab-1* 3′ enhancer segments showed greatest activity shortly after mid-day (ZT ∼10), when their enhancer strength was approximately 60–100% higher than at its lowest point (ZT ∼22). In contrast, the *rbcS-E9* 3′ enhancer segment was strongest close to midnight (ZT 19), and its enhancer strength dropped to approximately 40% of its maximum during the day (Figure 4B).

Next, we sought to identify mutations in the *AB80*, *Cab-1*, and *rbcS-E9* 3′ enhancer segments that disrupt their circadian regulation. We found that the amplitude of the circadian oscillation in enhancer strength remained largely unchanged across all single-nucleotide enhancer variants in our library (Figure 4C; Supplemental Figure S9– Supplemental Figure S12; Supplemental Data Set 2). Nearly all enhancer variants showed activity profiles across time points that matched their corresponding wild-type enhancer, especially when variants with very low and hence noisy enhancer strength values were excluded from the analysis (Figure 4D). Consistent with the observed lack of circadian cycle-specific variant effects, variant effects measured in this experiment correlated well with those measured in long-day light/dark cycles (Figure 4C; Supplemental Figure S8B). In summary, the *AB80*, *Cab-1*, and *rbcS-E9* enhancers are regulated by the circadian clock, and this regulation is robustly encoded in their sequence, as individual single-nucleotide mutations did not abolish it. The observed robustness of the circadian regulation is likely due to the presence of multiple binding sites for transcription factors controlled by the circadian clock in each enhancer. Further experiments are needed to test this hypothesis.

### Epistatic interactions between mutations in adjacent mutation-sensitive regions

Given the observed differences in activity caused by the same mutations when present in the 5’ vs. 3’ enhancer segment of the three enhancers, we hypothesized that the mutation-sensitive regions interact in a cooperative manner. To test this hypothesis, we looked for epistatic interactions between pairs of mutations in mutation-sensitive regions. In the absence of cooperativity, the change in enhancer strength relative to the wild-type enhancer caused by two mutations should equal the sum of the changes observed for the corresponding single mutations. In contrast, epistatic interactions between individual mutations would lead to less reduction in enhancer strength than expected based on the single mutation effects.

We created a Plant STARR-seq library containing single-nucleotide deletion variants of the *AB80* (19 deletions), *Cab-1* (32 deletions), and *rbcS-E9* (25 deletions) 3′ enhancer segments, as well as all possible variants containing a combination of two of these deletions within the same enhancer segment (Figure 5A). The enhancer strength of the single-nucleotide deletion variants in this small library correlated well with the values measured in the comprehensive single-nucleotide variant library (Supplemental Figure S13A).

**Figure 5.**
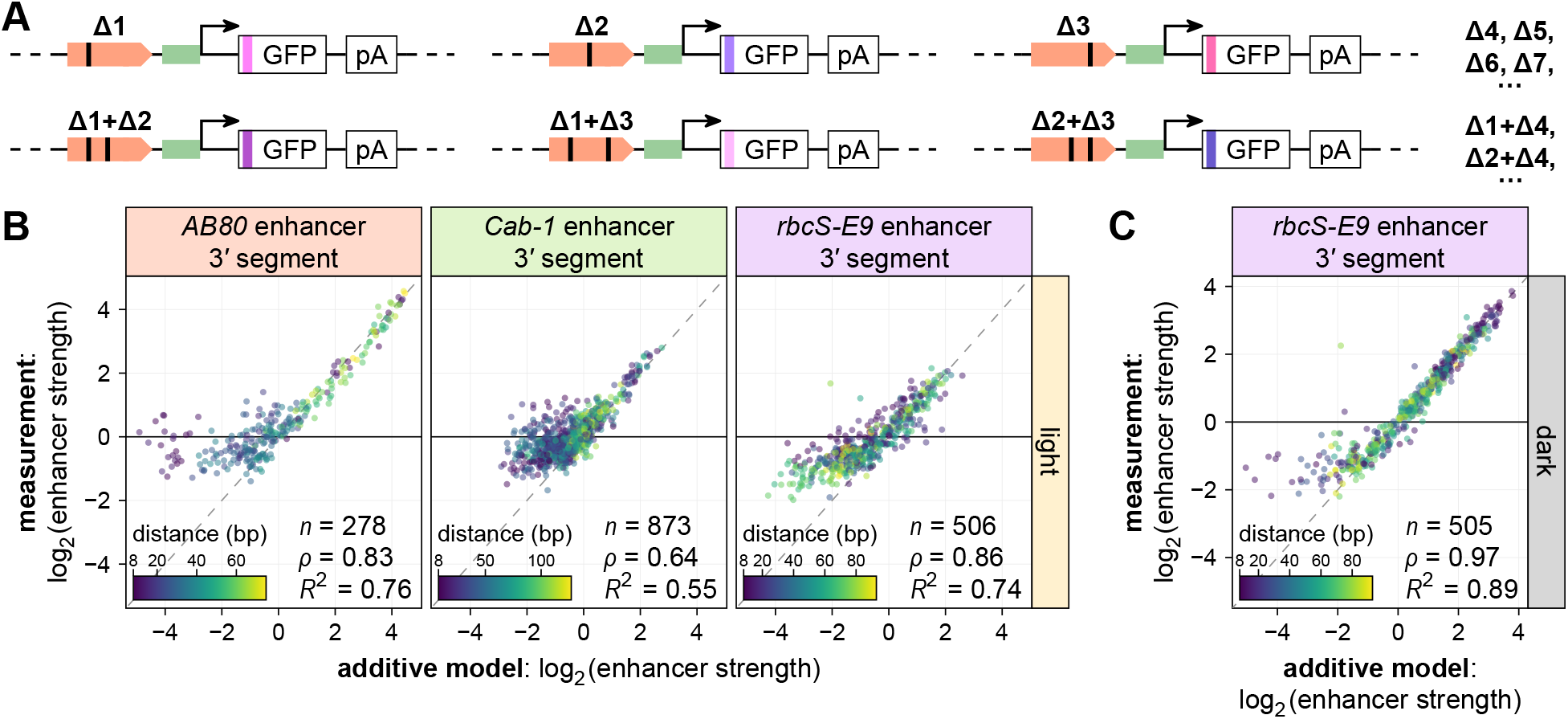
Epistatic interactions between single-nucleotide deletions. **A**–**C**, Selected single-nucleotide deletion variants (**A**; Δ1, Δ2, Δ3, …) of the 3′ segment of the *AB80*, *Cab-1*, and *rbcS-E9* enhancers and all possible combinations with two of these deletions (**A**; Δ1+Δ2, Δ1+Δ3, Δ2+Δ3, …) were subjected to Plant STARR-seq in tobacco plants grown in normal light/dark cycles (**B**) or completely in the dark (**C**) for two days prior to RNA extraction. For each pair of deletions, the expected enhancer strength based on the sum of the effects of the individual deletions (additive model) is plotted against the measured enhancer strength. The color of the points represents the distance between the two deletions in a pair.

We compared the strength of the enhancer variants carrying two deletions to predictions based on the additive strength of the two single deletions and found strong concordance overall (Figure 5B). When tested in the light, however, several variants with pairs of deletions showed greater enhancer strength than predicted (Figure 5B). In these cases, the two deletions tended to be relatively close to each other (8-20 bp; deletion pairs with a distance of less than 8 bp were excluded from this analysis). The same analysis for the *rbcS-E9* enhancer in the dark found fewer deletion pairs with epistatic interactions than in the light (Figure 5C). The strength of the *AB80* and *Cab-1* enhancers in the dark is too low to draw reliable conclusions from this analysis.

Taken together, we observed epistatic interactions between deletions in mutation-sensitive regions of the *AB80*, *Cab-1*, and *rbcS-E9* enhancers. The effect of these interactions dissipated over distance and was most pronounced in the light.

### The number, spacing, and order of mutation-sensitive regions affects enhancer strength

We selected 20 short (17–47 bp) fragments of the *AB80*, *Cab-1*, and *rbcS-E9* enhancers that span one to three mutation-sensitive regions to address how the number, spacing and order of mutation-sensitive regions affect enhancer strength. We also selected two control fragments, one derived from a mutation-insensitive region of the *Cab-1* enhancer and another from a randomly shuffled version of the *AB80* fragment d. Individual fragments as well as synthetic enhancers (combinations of two or three fragments) were cloned upstream of the 35S minimal promoter, and their enhancer strength in the light and dark was measured with Plant STARR-seq (Figure 6A; Supplemental Data Set 3). The individual fragments showed little to no enhancer activity in either condition (Figure 6B), consistent with our finding on cooperativity of mutation-sensitive regions. To systematically compare the enhancer strength of the individual enhancer fragments to the results of the saturation mutagenesis, we used the area under the curve in the mutational sensitivity plots (Figure 2) as a proxy for how much a fragment contributes to the strength of the full-length enhancer. In the light, there was no correlation between the strength of the enhancer fragments and their respective area under the curve observed in the saturation mutagenesis. In contrast, these metrics were well correlated in the dark (Figure 6C).

**Figure 6.**
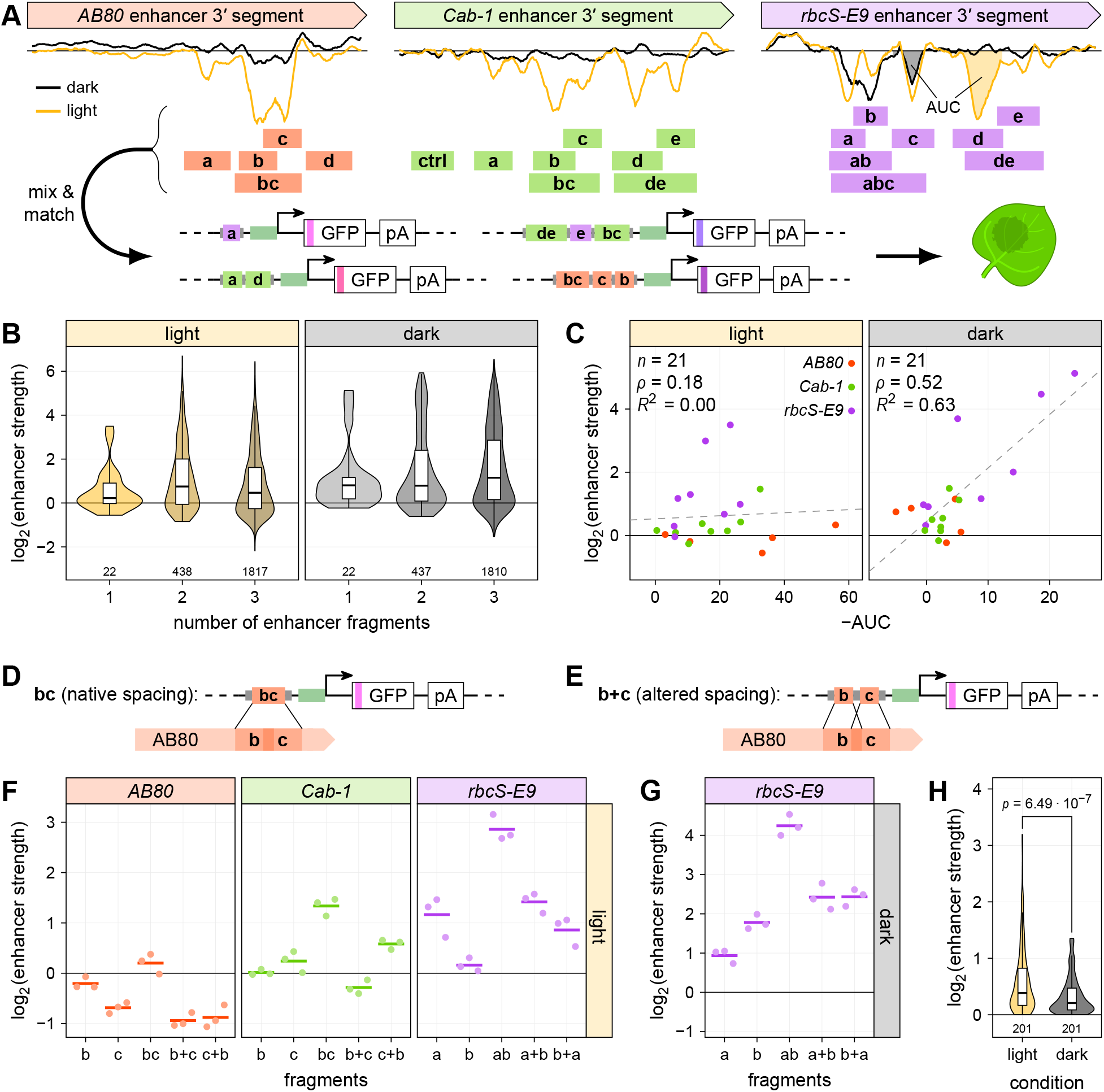
The number, spacing, and order of mutation-sensitive regions affects enhancer strength. **A**, Fragments of the *AB80*, *Cab-1*, and *rbcS-E9* enhancers spanning 1–3 mutation-sensitive regions (shaded rectangles; labeled a–e, ab, abc, bc, de) as well as a control fragment (ctrl) from a mutation-insensitive region in *Cab-1* and a shuffled version of the *AB80* fragment d were ordered as oligonucleotides. These fragments were randomly combined to create synthetic enhancers with up to three fragments which were then subjected to Plant STARR-seq in tobacco plants grown in normal light/dark cycles (light) or completely in the dark (dark) for two days prior to RNA extraction. The mutational sensitivity plots are reproduced from Figure 2. **B**, Violin plots of the strength of the synthetic enhancers grouped by the number of contained fragments. **C**, For each enhancer fragment, the area under the curve (AUC) in the mutational sensitivity plots was calculated and plotted against the fragment’s enhancer strength. AUCs in the dark or light for *rbcS-E9* fragments c and d, respectively, are shown in **A**. Pearson’s *R*^2^, Spearman’s ρ, and number (*n*) of enhancer fragments are indicated. A linear regression line is shown as a dashed line. **D**–**G**, Plots of the strength of enhancer fragments (**D**) or fragment combinations (separated by a + sign and shown in the order in which they appear in the construct; **E**) in three replicates (points) and the mean strength (lines). Enhancer strength was determined using tobacco plants grown in the light (**F**) or dark (**G**) prior to RNA extraction. **H**, Violin plots of the difference in enhancer strength between synthetic enhancers harboring the same two enhancer fragments but in different order. The *p*-value from a two-sided Wilcoxon rank-sum test comparing light and dark results is indicated (*p*). Violin plots in **B** and **H** represent the kernel density distribution and the box plots inside represent the median (center line), upper and lower quartiles and 1.5× interquartile range (whiskers) for all corresponding synthetic enhancers. Numbers at the bottom of each violin indicate the number of elements in each group. Enhancer strength in **B**–**G** was normalized to a control construct without an enhancer (log_2_ set to 0).

Most synthetic enhancers showed only weak enhancer activity (Figure 6B), indicating that simply combining mutation-sensitive regions is not sufficient for cooperative interactions. To test if the spacing of mutation-sensitive regions affects enhancer activity, we compared the strength of enhancer fragments spanning two such regions (with both regions present at the same distance as in the wild-type enhancer; named ab, bc, de; Figure 6D) to synthetic enhancers composed of the two individual fragments (with altered spacing between the mutation-sensitive regions; named a+b, b+c, d+e; Figure 6E). In almost all cases, the fragment spanning two mutation-sensitive regions was stronger than the combination of the individual fragments (Figure 6, F and G; Supplemental Figure S14). In addition to the spacing, the order of enhancer fragments also affected the strength of the resulting synthetic enhancers, and this effect was stronger in the light (Figure 6, F–H).

In summary, the activity of the full-length enhancers is the result of cooperative interactions between their constituent mutation-sensitive regions, in particular in the light. This interpretation is supported by the observation that the enhancer strength of the combinations of mutation-sensitive regions depends on their spacing and order. In contrast, the mutation-sensitive regions of the *rbcS-E9* enhancer function largely independently and additively in the dark. Because we analyzed only three plant enhancers, it remains to be tested how generalizable our findings are.

### Enhancer fragments can be used to design synthetic enhancers

Although many of the synthetic enhancers created by combining fragments of the *AB80*, *Cab-1*, and *rbcS-E9* enhancers were inactive (940 enhancers), more than half of them showed activity in at least one condition (1,364 enhancers with log_2_[enhancer strength] > 1; Figure 7A). Of the active synthetic enhancers, most showed highest strength in the dark (728 enhancers). In contrast, only 203 synthetic enhancers were active specifically in the light, despite being comprised largely of mutation-sensitive regions found in the light. Finally, 433 synthetic enhancers were active in both the light and the dark. We validated these results by retesting a subset of approximately 400 synthetic enhancers in a separate library (Supplemental Figure S13B). Moreover, we measured the activity of 11 synthetic enhancers in stable *Arabidopsis* lines using the dual-luciferase assay (Figure 7B) and observed strong correlation with the enhancer strengths measured by Plant STARR-seq (Figure 7C).

**Figure 7.**
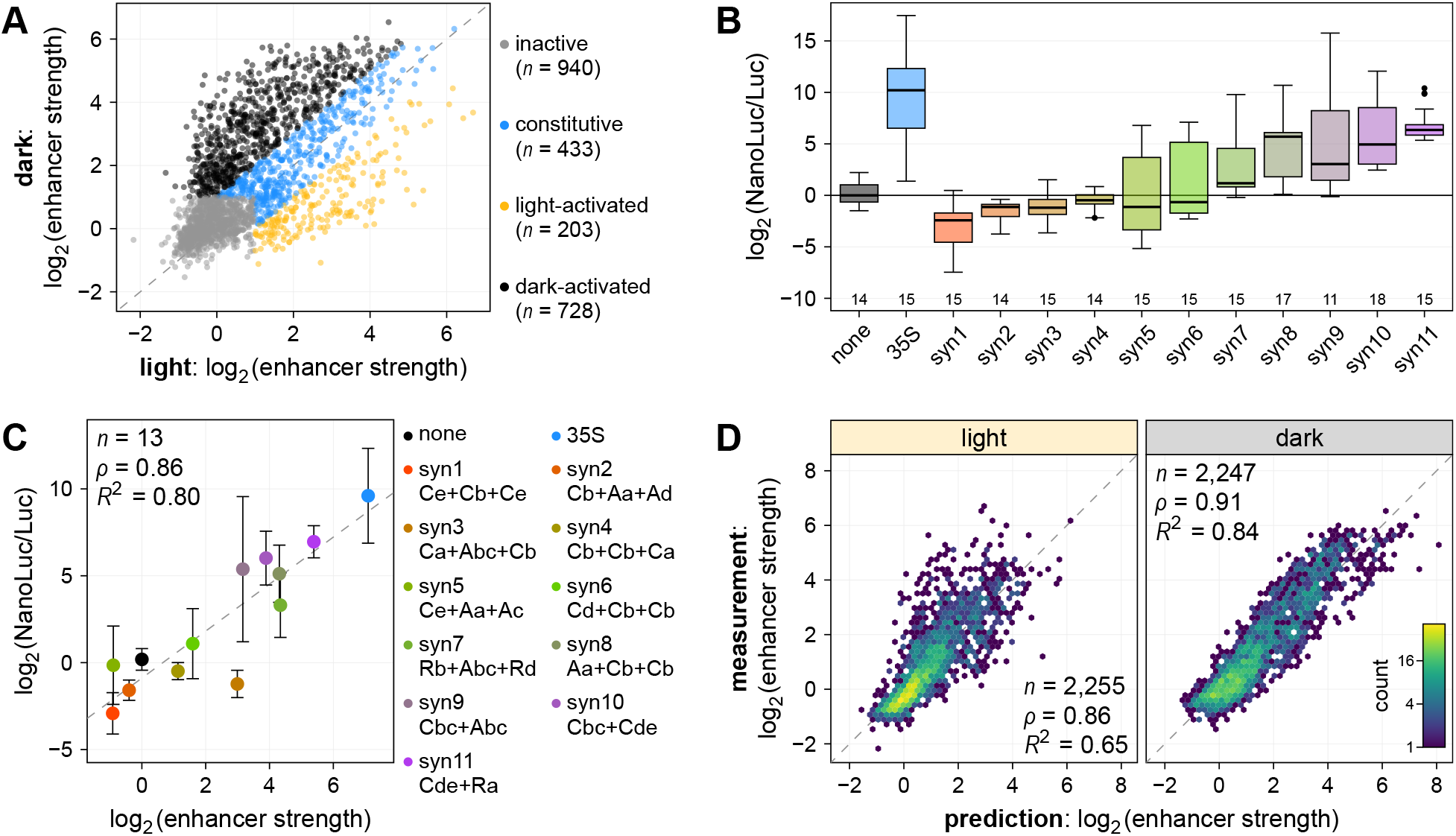
Enhancer fragments can be used to build condition-specific synthetic enhancers. **A**, Plot of the strength of synthetic enhancers created by randomly combining up to three fragments derived from mutation-sensitive regions of the *AB80*, *Cab-1*, and *rbcS-E9* enhancers (see Figure 6A) as measured by Plant STARR-seq in the light or dark. The synthetic enhancers were grouped into four categories: inactive, log_2_(enhancer strength) ≤ 1 in both conditions; constitutive, similar strength in both conditions; light-activated, at least two-fold more active in the light; dark-activated, at least two-fold more active in the dark. The number (*n*) of synthetic enhancers in each category is indicated. **B**, Dual-luciferase reporter constructs (see Figure 1D) were created for 11 synthetic enhancers (syn1–11). Nanoluciferase activity was measured in at least 4 T2 plants from these lines and normalized to the activity of luciferase. The NanoLuc/Luc ratio was normalized to a control construct without an enhancer (none; log_2_ set to 0). Box plots are as defined in Figure 1E. **C**, The mean NanoLuc/Luc ratio was compared to the mean enhancer strength determined by STARR-seq. A linear regression line is shown as a dashed line. Error bars represent the 95% confidence interval. The constituent fragments of the synthetic enhancers are indicated with fragments separated by a + sign. The first letter indicates the enhancer from which the fragment is derived (A, *AB80*; C, *Cab-1*; R, *rbcS-E9*) and the lowercase letters represent the fragment name. **D**, A linear model was built to predict the strength of the synthetic enhancers based on the strength of the constituent individual fragments. Hexbin plots (color represents the count of points in each hexagon) of the correlation between the model’s prediction and the measured data are shown. In **C** and **D**, Pearson’s *R*^2^, Spearman’s ρ, and number (*n*) of synthetic enhancers are indicated.

When analyzing the composition of the synthetic enhancers, we found that dark-activated and constitutive synthetic enhancers often contained fragments of *rbcS-E9*, while inactive and light-activated synthetic enhancers often contained fragments of *AB80* and *Cab-1*. Many (57%) dark-activated synthetic enhancers contained at least one copy of *rbcS-E9* region b. Light-activated synthetic enhancers tended to contain *Cab-1* regions b – e and *AB80* regions b and c, but without clear preference for either region. Finally, constitutive synthetic enhancers commonly contained *rbcS-E9* regions a – b. These results are consistent with the condition-specific mutational sensitivity of these enhancers and enhancer regions.

Lastly, we asked if the strength of a given synthetic enhancer can be predicted based on the strength of its constituent fragments, despite the observed cooperativity in the light. A simple linear model based on the enhancer strength of the 22 individual enhancer fragments was able to predict the strength of synthetic enhancers with high accuracy (Figure 7D). Consistent with our cooperativity results, the model performed best when applied to measurements obtained in the dark (*R*^2^ = 0.84), but it also showed predictive power in the light (*R*^2^ = 0.65). A similar model that predicted the light-responsiveness of the synthetic enhancers showed somewhat lower accuracy (*R^2^* = 0.42; Supplemental Figure S15). In short, a relatively small number of well-characterized enhancer fragments is sufficient to create a diverse set of short (less than 150 bp) synthetic enhancers with condition-specific activity.

## Discussion

In this study, we used Plant STARR-seq to characterize the *AB80*, *Cab-1*, and *rbcS-E9* enhancers under two different light regimes. We observed that, in the light, the activity of the three enhancers is influenced by their grammar and by cooperative interactions between constituent functional regions defined by mutational sensitivity. Thus, these plant enhancers appear to show enhanceosome features in the light. Enhanceosomes in animal cells contain dense arrays of transcription factor binding sites in specific orientation, spacing, and order that together recruit transcription factor complexes (Thanos and Maniatis, 1995; Panne, 2008); full activity requires cooperative binding of all complex members. In the dark, the three plant enhancers show stronger resemblance to billboard or transcription factor collective enhancers, with reduced dependency on grammar and cooperativity.

Because of the small number of plant enhancers tested, we cannot conclude whether enhancer activity in the light generally relies more on cooperativity than activity in the dark. It is tempting, however, to speculate that enhancers that respond to stimuli may adhere more to an enhanceosome model because they need to integrate multiple environmental signals. This integration, which is critical for plant growth and development, would be expected to involve multiple different transcription factors acting in concert. However, we presently lack such transcription factor binding data in our assay.

We identify regions important for enhancer activity and demonstrate that these regions harbor putative binding sites for transcription factors. A previous study identified several regions of the *Cab-1* enhancer that are protected from DNase I digestion in the presence of a nuclear extract from light-grown tobacco (Gotor et al., 1993). These DNase I-protected regions overlap with the mutation-sensitive regions of *Cab-1* identified here, supporting the hypothesis that mutation-sensitive regions are bound by transcription factors *in planta*.

As transcription factors of the same family often bind to highly similar motifs (O’Malley et al., 2016; Jores et al., 2021; Zenker et al., 2023), we cannot pinpoint the specific family member(s) binding to a given site. Nonetheless, our results point to an important role of MYB family transcription factors in generating enhancer activity of the *AB80*, *Cab-1*, and *rbcS-E9* enhancers in the light and in the dark. While it might seem counterintuitive that the same transcription factors are recruited to enhancers that function in different conditions (light or dark) and via different models (enhanceosome or billboard), similar observations have been made in *Drosophila melanogaster* (Liu and Posakony, 2012).

Our results highlight the crucial role that enhancer grammar can play in determining enhancer strength. However, future studies will be required to decipher in-depth the rules of how the order, spacing and orientation of transcription factor binding sites affects activity or if such rules even exist. This knowledge is crucial for the rational *de novo* design of highly specific enhanceosome enhancers, which is not possible to date. As an alternative, we demonstrate that enhancer fragments with known activity can be combined in an additive, billboard-like fashion to build synthetic enhancers with high activity and condition-specificity. The success of this approach is rooted in the hierarchical structure of enhancers (Fromental et al., 1988; Ondek et al., 1988). At the base level, enhancers consist of transcription factor binding sites (called “enhansons” in the 1988 studies) that are combined with an optimized grammar to form short “proto-enhancers” or “enhancer elements.” These proto-enhancers are then combined with a more flexible grammar to form full-length enhancers.

This study showcases the power of Plant STARR-seq to characterize enhancers at nucleotide resolution. A similar approach could be used to characterize other *cis*-regulatory elements such as promoters, silencers, or insulators. However, the design and interpretation of Plant STARR-seq experiments has limitations. First, Plant STARR-seq is best suited for highly efficient plant transformation systems that enable high coverage of the tested sequences. Thus far, this efficiency is possible only in transient assays in a few species such as tobacco and maize, and only in a few tissues (Jores et al., 2020, 2021). As implemented currently, Plant STARR-seq misses most tissue- and species-specific effects, and it cannot replicate the endogenous genome and chromatin context. Second, Plant STARR-seq relies on array synthesis to generate large libraries of candidate regulatory elements. Because accurate array synthesis is currently limited to short sequences, Plant STARR-seq cannot detect long-range regulatory interactions. These two limitations can be overcome by improved plant transformation protocols and improved technologies for the accurate synthesis or assembly of long test sequences. Furthermore, improved methods for plant genome engineering will enable researchers to validate their findings in the context of native plant genomes.

## Material and methods

### Library design and construction

The full-length *AB80*, *Cab-1*, and *rbcS-E9* enhancers and their 169-bp 3′ and 5′ segments as well as the 35S enhancer were PCR amplified from pZS*11_4enh (Addgene no. 149423; https://www.addgene.org/149423/; Jores et al., 2020). The sequences of all oligonucleotides used for cloning and sequencing are included in Supplemental Data Set 4.

Single-nucleotide and double-deletion variants of the three plant enhancers with 15-bp flanking sequences (5′ TAACTCGCCTCGATC and 3′ CCGTGACAGGTCATT for *AB80*; 5′ ACATGGGGGATCATG and 3′ GCATTAGCCAGTCTG for *Cab-1*; 5′ GCTCCAGTTCCCAAC and 3′ GATCGTCCAGTCTGA for *rbcS-E9*) for amplification were ordered as an oligonucleotide array from Twist Bioscience.

Enhancers fragments (Figure 6A) with 5′ GTGATG overhangs and their reverse-complements with 5′ CATCAC overhangs were ordered as oligonucleotides, annealed, and 5′ phosphorylated with T4 Polynucleotide Kinase (NEB). The fragments were mixed with Golden Gate cloning adaptors (5′ adapter: GAGAGGGTCTCCACTC and CATCACGAGTGGAGACCCTCTC; 3′ adapter: GTGATGAGGACGAGACCCTCTC and GAGAGGGTCTCGTCCT; annealed and 5′ phosphorylated) and ligated with T4 DNA ligase. The ligation products were size-selected to exclude constructs over 150 bp. Enhancer fragment combinations for the validation library were ordered as an oligonucleotide array from Twist Bioscience.

All libraries used in this study were constructed using pPSup (Addgene no. 149416; https://www.addgene.org/149416/; Jores et al., 2020) as the base plasmid. The plasmid’s T-DNA region harbors a phosphinothricin resistance gene (BlpR) and a GFP reporter construct terminated by the poly(A) site of the *Arabidopsis thaliana* ribulose bisphosphate carboxylase small chain 1A gene. The 35S minimal promoter followed by the synthetic 5′ UTR synJ (ACACGCTGGAATTCTAGTATACTAAACC; Kanoria and Burma, 2012), an ATG start codon and a 15-bp random barcode (VNNVNNVNNVNNVNN; V = A, C, or G) was cloned in front of the second codon of GFP by Golden Gate cloning (Engler et al., 2008) using BbsI-HF (NEB). Enhancers, enhancer variants, and enhancer fragment combinations were cloned upstream of the 35S minimal promoter by Golden Gate cloning using BsaI-HFv2 (NEB). The resulting libraries were bottlenecked to yield 10–20 barcodes per enhancer.

The base plasmid pDL for dual-luciferase constructs was created from pPSup after the 35S minimal promoter and synJ 5′ UTR were inserted into the BbsI Golden Gate site. The GFP reporter gene was replaced with nanoluciferase by Gibson Assembly. A second round of Gibson Assembly was used to insert the luciferase gene driven by the *Arabidopsis UBQ10* promoter (TAIR10 Chr4:2716532–2718558) and terminated by the 35S terminator between the T-DNA left border and the BlpR gene. The nanoluciferase coding sequence and the 35S terminator were ordered as synthesized DNA fragments. The luciferase coding sequence was amplified from pGreen_dualluc_3’UTR_sensor (Addgene no. 55206; https://www.addgene.org/55206/; Liu et al., 2014) and the *UBQ10* promoter was amplified for *Arabidopsis* Col-0 genomic DNA. The assembled plasmid pDL was deposited at Addgene (Addgene no. 208978; https://www.addgene.org/208978/). The 3′ segments of the *AB80*, *Cab-1*, and *rbcS-E9* enhancers and the synthetic enhancers syn1–11 were cloned upstream of the 35S minimal promoter in pDL by Golden Gate cloning using BsaI-HFv2 (NEB). The synthetic enhancers were ordered as synthesized DNA fragments.

### Plant cultivation and transformation

Tobacco (*Nicotiana benthamiana*) was grown in soil (Sunshine Mix no. 4) at 25°C in a long-day photoperiod (16 h light and 8 h dark; cool-white fluorescent lights [Philips TL-D 58 W/840]; intensity 300LμmolLm^-2^Ls^-1^). Plants were transformed approximately 3Lweeks after germination. For transient transformation of tobacco leaves, enhancer libraries were introduced into *Agrobacterium tumefaciens* strain GV3101 (harboring the virulence plasmid pMP90 and the helper plasmid pSoup) by electroporation. An overnight culture of the transformed *A. tumefaciens* was diluted into 100Lml YEP medium (1% [w/v] yeast extract and 2% [w/v] peptone) and grown at 28°C for 8 h. A 5-ml input sample of the cells was collected, and plasmids were isolated from it using the QIAprep Spin Miniprep Kit (QIAGEN) according to the manufacturer’s instructions. The remaining cells were harvested and resuspended in 100 ml induction medium (M9 medium [3 g/L KH_2_PO_4_, 0.5 g/L NaCl, 6.8 g/L Na_2_HPO_4_, and 1 g/L NH_4_Cl] supplemented with 1% [w/v] glucose, 10 mM MES, pHL5.2, 100 μM CaCl_2_, 2LmM MgSO_4_, and 100LμM acetosyringone). After overnight growth, the *Agrobacteria* were harvested, resuspended in infiltration solution (10 mM MES, pHL5.2, 10LmM MgCl_2_, 150 μM acetosyringone, and 5 μM lipoic acid) to an optical density of 1 and infiltrated into leaves 3 and 4 of six tobacco plants. The plants were further grown for 48Lh under normal conditions (16 h light and 8 h dark) or in the dark before mRNA extraction.

*Arabidopsis thaliana* Col-0 was grown in soil (Sunshine Mix no. 4) at 20°C in a long-day photoperiod (16 h light and 8 h dark; cool-white fluorescent lights [Sylvania FO32/841/ECO 32W]; intensity 100LμmolLm^-2^Ls^-1^). For transformation, dual-luciferase plasmids were introduced into *Agrobacterium tumefaciens* strain GV3101 (harboring the virulence plasmid pMP90 and the helper plasmid pSoup) by electroporation. Transgenic *Arabidopsis* plants were generated by floral dipping (Clough and Bent, 1998) and selected for by spraying with a 0.01% Glufosinate solution.

### Plant STARR-seq

For all Plant STARR-seq experiments, at least two independent biological replicates were performed. Different plants and fresh *Agrobacterium* cultures were used for each biological replicate.

Transiently transformed tobacco leaves were harvested 2 days after infiltration and partitioned into two batches of 6 leaves each. The leaf batches were frozen in liquid nitrogen, finely ground with mortar and pestle, and immediately resuspended in 12 mL TRIzol (Thermo Fisher Scientific). The suspensions from both leaf batches were pooled and cleared by centrifugation (5 min, 4,000 x g, 4°C). The supernatant was mixed with 5 mL chloroform and centrifuged (15 min, 4,000 x g, 4°C). The upper, aqueous phase was transferred to a new tube and was washed once more with 5 mL chloroform. The aqueous phase (approximately 10 mL) was transferred to a new tube, and mixed by inversion with 10 mL isopropanol and 10 mL high salt buffer (0.8 M sodium citrate, 1.2 M NaCl). The solution was incubated for 15 min at RT to precipitate the RNA and centrifuged (30 min, 4,000 x g, 4°C). The pellet was washed in 25 mL ice-cold 70% ethanol, centrifuged (5 min, 4000 x g, 4°C), and air-dried. The pellet was resuspended in 2.4 mL of warm (65°C) nuclease-free water and split into two aliquots. From each aliquot, mRNAs were isolated using 150 µL magnetic Oligo(dT)_25_ beads (Thermo Fisher Scientific) according to the manufacturer’s instructions. Elution was performed with 40 µL 10 mM Tris (pH 7.4) and the eluates from both aliquots were pooled. To remove DNA contaminations, the mRNA solution was mixed 10 μL DNase I buffer without MnCl_2_, 10 μL 100 mM MnCl_2_, 1 μL RNaseOUT, and 2 μL DNase I (Thermo Fisher Scientific), and incubated for 1 h at 37°C. To precipitate the mRNA, 1 μL 20 mg/mL glycogen (Thermo Fisher Scientific), 10 µL ice-cold 8M LiCl and 250 µL ice-cold 100% ethanol was added. After incubation for 15 min at −80°C, the RNA was pelleted by centrifugation (20 min, 20,000 x g, 4°C). The pellet was washed with 200 µL ice-cold 70% ethanol, centrifuged (5 min, 20,000 x g, 4°C), air-dried, and resuspended in 100 µL nuclease-free water. For cDNA synthesis, eight reactions with 11 µL mRNA solution, 1 µL 2 µM GFP-specific reverse transcription primer, and 1 µL 10 mM dNTPs were incubated at 65°C for 5 min then immediately placed on ice. The reactions were supplemented with 4 µL 5X SuperScript IV buffer, 1 µL 100 mM DTT, 1 µL RNaseOUT, and 1 µL SuperScript IV reverse transcriptase (ThermoFisher Scientific). To ensure that the samples were largely free of DNA contamination, four reactions were used as controls, where the reverse transcriptase and RNaseOUT were replaced with water. Reactions were incubated for 10 min at 55°C, followed by 10 min at 80°C. Sets of 4 reactions each were pooled. The cDNA was purified with the Zymo Clean&Concentrate-5 kit, and eluted in 20 µL 10 mM Tris. The barcode was amplified with 10-20 cycles of polymerase chain reaction (PCR) and read out by next generation sequencing.

### Subassembly and barcode sequencing

Paired-end sequencing on an Illumina NextSeq 550 platform was used to link enhancers to their respective barcodes. The enhancer region was sequenced using paired 144-bp reads, and two 15-bp indexing reads were used to sequence the barcodes. The paired enhancer and barcode reads were assembled using PANDAseq (version 2.11; Masella et al., 2012). Enhancer-barcode pairs with less than 5 reads and enhancers with a mutation or truncation were discarded.

For each Plant STARR-seq experiment, barcodes were sequenced using paired-end reads on an Illumina NextSeq 500, 550, or 2000 system. The paired barcode reads were assembled using PANDAseq.

### Computational methods

The code used for the analysis and to generate the figures is available on GitHub (https://github.com/tobjores/cooperativity-and-additivity-in-plant-enhancers).

For analysis of the Plant STARR-seq experiments, the reads for each barcode were counted in the input and cDNA samples. Barcode counts below 5 and barcodes present in only one replicate were discarded. Barcode counts were normalized to the sum of all counts in the respective sample. For barcodes, enhancer strength was calculated by dividing the normalized barcode counts in the cDNA sample by that in the corresponding input sample. The sum of the normalized counts for all barcodes associated with a given enhancer, enhancer variant, or enhancer fragment combination were used to calculate its strength. For each replicate, enhancer strength was normalized to the median enhancer strength. Unless indicated otherwise, the mean enhancer strength across all replicates was used for analyses. Spearman and Pearson’s correlation were calculated using base R (version 4.3.1).

To identify putative transcription factor binding sites within the *AB80*, *Cab-1*, and *rbcS-E9* enhancers, we used the universalmotif package (version 1.18.1) in R to scan the enhancer sequences for matches on each strand to known transcription factor binding motifs (obtained from Jores et al., 2021). Scores for how well a variant sequence matches a transcription factor binding motif were calculated using the score_match() function of the universalmotif package. For the generation of sequence logo plots for mutation-sensitive regions we adapted previously published approaches (Andrilenas et al., 2018; Ireland et al., 2020). The enhancer strength of all variants within a given mutation-sensitive region was normalized to the strength of the wild-type variant, scaled by a factor β, and turned into an information content matrix using the create_motif() function of the universalmotif package. The scaling factor β was chosen such that the final motif had an average per-base information content of 1. The generated motifs were compared to known transcription factor binding motifs using the compare_motifs() function of the universalmotif package.

For the generation of circadian rhythm curves, a linear model was fitted to the time course Plant STARR-seq data using the lm() function in R with the formula: log2(enhancer strength) = sin(2 · π · time / 24) + cos(2 · π · time / 24), where time refers to the time in hours when the samples were harvested.

To predict the strength of synthetic enhancers generated by combining enhancer fragments, a liner model was fitted to Plant STARR-seq data using the lm() function in R with the formula: log2(enhancer strength) = log2(enhancer strength fragment 3) + log2(enhancer strength fragment 2) + log2(enhancer strength fragment 1), where log2(enhancer strength fragment 1–3) is the enhancer strength of the corresponding fragment when tested individually. Fragments are numbered by increasing distance from the minimal promoter (fragment 1 is the fragment closest to the promoter, fragment 3 the most distal one). For combinations of two fragments, log2(enhancer strength fragment 3) was set to 0.

### Dual-luciferase assay

Transgenic *Arabidopsis* lines (T2 generation) with dual-luciferase constructs were grown in soil for 3 weeks. A cork borer (4 mm diameter) was used to collect a total of 3 leaf discs from the third and fourth leaf of the plants. The leaf discs were transferred to 1.5 mL tubes filled with approximately 10 glass beads (1 mm diameter), snap-frozen in liquid nitrogen, and disrupted by shaking twice for 5 sec in a Silamat S6 (Ivoclar) homogenizer. The leaf disc debris was resuspended in 100 µL 1X Passive Lysis Buffer (Promega). The solution was cleared by centrifugation (5 min, 20,000 x g, RT) and 10 µL of the supernatant were mixed with 90 µL 1X passive lysis buffer. Luciferase and nanoluciferase activity were measured on a Biotek Synergy H1 plate reader using the Promega Nano-Glo Dual-Luciferase Reporter Assay System according to the manufacturer’s instructions. Specifically, 10 µL of the leaf extracts were combined with 75 µL ONE-Glo EX Reagent, mixed for 3 min at 425 rpm, and incubated for 2 min before measuring luciferase activity. Subsequently, 75 µL NanoDLR Stop&Glo Reagent were added to them sample. After 3 min mixing at 425 rpm and 12 min incubation, nanoluciferase activity was measured. Three independent biological replicates were performed.

### Accession Numbers

The code used for the analysis and to generate the figures is available on GitHub (https://github.com/tobjores/cooperativity-and-additivity-in-plant-enhancers). All barcode sequencing reads were deposited in the National Center for Biotechnology Information (NCBI) Sequence Read Archive under the BioProject accession PRJNA1015372 (http://www.ncbi.nlm.nih.gov/bioproject/1015372/). The sequences for the enhancers can be obtained from NCBI (https://www.ncbi.nlm.nih.gov/nuccore/) using the following accessions: V00141 (35S), X03074 (AB80), X05823 (Cab-1), and X00806 (rbcS-E9).

## Supporting information

Supplemental Data Set 1

Supplemental Data Set 2

Supplemental Data Set 3

Supplemental Data Set 4

## Acknowledgements

This work was supported by the National Science Foundation (RESEARCH-PGR grant no. 1748843 to S.F. and C.Q. and PlantSynBio grant no. 2240888 to C.Q.), the German Research Foundation (DFG; postdoctoral fellowship no. 441540116 to T.J., and Emmy Noether programme grant no. 517938232 to T.J.), the National Institutes of Health (T32 training grant no. HG000035 to J.T., NIGMS grant no. R01-GM079712 to J.T.C. and C.Q., and NIGMS MIRA grant no. 1R35GM139532 to C.Q.), and the United States Department of Agriculture (NIFA postdoctoral fellowship no. 2023-67012-39445 to N.A.M).

## Author contributions

All authors conceived and interpreted experiments; T.J., J.T., and N.A.M performed experiments; T.J., and J.T. analyzed the data; T.J., S.F., and C.Q. prepared the manuscript and figures. All authors read and revised the manuscript.

## References

Andrilenas KK, Ramlall V, Kurland J, Leung B, Harbaugh AG, Siggers T. DNA-binding landscape of IRF3, IRF5 and IRF7 dimers: implications for dimer-specific gene regulation. Nucleic Acids Res. 2018:46(5):2509–2520. 10.1093/nar/gky002

Arnosti DN, Kulkarni MM. Transcriptional enhancers: Intelligent enhanceosomes or flexible billboards? J. Cell. Biochem. 2005:94(5):890–898. 10.1002/jcb.20352

Banerji J, Rusconi S, Schaffner W. Expression of a β-globin gene is enhanced by remote SV40 DNA sequences. Cell. 1981:27(2, Part 1):299–308. 10.1016/0092-8674(81)90413-X

Benfey PN, Ren L, Chua NH. Tissue-specific expression from CaMV 35S enhancer subdomains in early stages of plant development. EMBO J. 1990:9(6):1677–1684.

de Boer CG, Vaishnav ED, Sadeh R, Abeyta EL, Friedman N, Regev A. Deciphering eukaryotic gene-regulatory logic with 100 million random promoters. Nat. Biotechnol. 2020:38(1):56–65. 10.1038/s41587-019-0315-8

Cai Y-M, Kallam K, Tidd H, Gendarini G, Salzman A, Patron NJ. Rational design of minimal synthetic promoters for plants. Nucleic Acids Res. 2020:48(21):11845–11856. 10.1093/nar/gkaa682

Carré IA, Kim J. MYB transcription factors in the Arabidopsis circadian clock. J. Exp. Bot. 2002:53(374):1551–1557. 10.1093/jxb/erf027

Clough SJ, Bent AF. Floral dip: a simplified method for Agrobacterium -mediated transformation of Arabidopsis thaliana. Plant J. 1998:16(6):735–743. 10.1046/j.1365-313x.1998.00343.x

Engler C, Kandzia R, Marillonnet S. A One Pot, One Step, Precision Cloning Method with High Throughput Capability. PLOS ONE. 2008:3(11):e3647. 10.1371/journal.pone.0003647

Erhard KF Jr, Talbot J-ERB, Deans NC, McClish AE, Hollick JB. Nascent Transcription Affected by RNA Polymerase IV in Zea mays. Genetics. 2015:199(4):1107–1125. 10.1534/genetics.115.174714

Fang RX, Nagy F, Sivasubramaniam S, Chua NH. Multiple cis regulatory elements for maximal expression of the cauliflower mosaic virus 35S promoter in transgenic plants. Plant Cell. 1989:1(1):141–150. 10.1105/tpc.1.1.141

Farley EK, Olson KM, Zhang W, Brandt AJ, Rokhsar DS, Levine MS. Suboptimization of developmental enhancers. Science. 2015:350(6258):325–328. 10.1126/science.aac6948

Farley EK, Olson KM, Zhang W, Rokhsar DS, Levine MS. Syntax compensates for poor binding sites to encode tissue specificity of developmental enhancers. Proc. Natl. Acad. Sci. 2016:113(23):6508–6513. 10.1073/pnas.1605085113

Fejes E, Pay A, Kanevsky I, Szell M, Adam E, Kay S, Nagy F. A 268 bp upstream sequence mediates the circadian clock-regulated transcription of the wheat Cab-1 gene in transgenic plants. Plant Mol. Biol. 1990:15(6):921–932. 10.1007/bf00039431

Fluhr R, Kuhlemeier C, Nagy F, Chua N-H. Organ-Specific and Light-Induced Expression of Plant Genes. Science. 1986:232(4754):1106–1112. 10.1126/science.232.4754.1106

Friedman RZ, Ramu A, Lichtarge S, Myers CA, Granas DM, Gause M, Corbo JC, Cohen BA, White MA. Active learning of enhancer and silencer regulatory grammar in photoreceptors. bioRxiv. 2023. 10.1101/2023.08.21.554146

Fromental C, Kanno M, Nomiyama H, Chambon P. Cooperativity and hierarchical levels of functional organization in the SV40 enhancer. Cell. 1988:54(7):943–953. 10.1016/0092-8674(88)90109-2

Gnesutta N, Chiara M, Bernardini A, Balestra M, Horner DS, Mantovani R. The Plant NF-Y DNA Matrix In Vitro and In Vivo. Plants. 2019:8(10):406. 10.3390/plants8100406

Gnesutta N, Kumimoto RW, Swain S, Chiara M, Siriwardana C, Horner DS, Holt BF III, Mantovani R. CONSTANS Imparts DNA Sequence Specificity to the Histone Fold NF-YB/NF-YC Dimer. Plant Cell. 2017:29(6):1516–1532. 10.1105/tpc.16.00864

Gotor C, Romero LC, Inouye K, Lam E. Analysis of three tissue-specific elements from the wheat Cab-1 enhancer. Plant J. 1993:3(4):509–518. 10.1046/j.1365-313X.1993.03040509.x

Hetzel J, Duttke SH, Benner C, Chory J. Nascent RNA sequencing reveals distinct features in plant transcription. Proc. Natl. Acad. Sci. 2016:113(43):12316–12321. 10.1073/pnas.1603217113

Ireland WT, Beeler SM, Flores-Bautista E, McCarty NS, Röschinger T, Belliveau NM, Sweredoski MJ, Moradian A, Kinney JB, Phillips R. Deciphering the regulatory genome of Escherichia coli, one hundred promoters at a time. eLife. 2020:9e55308. 10.7554/eLife.55308

Jindal GA, Farley EK. Enhancer grammar in development, evolution, and disease: dependencies and interplay. Dev. Cell. 2021:56(5):575–587. 10.1016/j.devcel.2021.02.016

Jores T, Hamm M, Cuperus JT, Queitsch C. Frontiers and techniques in plant gene regulation. Curr. Opin. Plant Biol. 2023:75102403. 10.1016/j.pbi.2023.102403

Jores T, Tonnies J, Dorrity MW, Cuperus JT, Fields S, Queitsch C. Identification of Plant Enhancers and Their Constituent Elements by STARR-seq in Tobacco Leaves. Plant Cell. 2020:32(7):2120–2131. 10.1105/tpc.20.00155

Jores T, Tonnies J, Wrightsman T, Buckler ES, Cuperus JT, Fields S, Queitsch C. Synthetic promoter designs enabled by a comprehensive analysis of plant core promoters. Nat. Plants. 2021:7(6):842–855. 10.1038/s41477-021-00932-y

Junion G, Spivakov M, Girardot C, Braun M, Gustafson EH, Birney E, Furlong EEM. A Transcription Factor Collective Defines Cardiac Cell Fate and Reflects Lineage History. Cell. 2012:148(3):473–486. 10.1016/j.cell.2012.01.030

Kanoria S, Burma PK. A 28 nt long synthetic 5′UTR (synJ) as an enhancer of transgene expression in dicotyledonous plants. BMC Biotechnol. 2012:12(1):85. 10.1186/1472-6750-12-85

Kim YJ, Rhee K, Liu J, Jeammet S, Turner MA, Small SJ, Garcia HG. Predictive modeling reveals that higher-order cooperativity drives transcriptional repression in a synthetic developmental enhancer. eLife. 2022:11e73395. 10.7554/eLife.73395

Kim S, Wysocka J. Deciphering the multi-scale, quantitative cis-regulatory code. Mol. Cell. 2023:83(3):373–392. 10.1016/j.molcel.2022.12.032

Kulkarni MM, Arnosti DN. Information display by transcriptional enhancers. Development. 2003:130(26):6569–6575. 10.1242/dev.00890

Laosuntisuk K, Elorriaga E, Doherty CJ. The Game of Timing: Circadian Rhythms Intersect with Changing Environments. Annu. Rev. Plant Biol. 2023:74(1):511–538. 10.1146/annurev-arplant-070522-065329

Liu F, Posakony JW. Role of Architecture in the Function and Specificity of Two Notch-Regulated Transcriptional Enhancer Modules. PLOS Genet. 2012:8(7):e1002796. 10.1371/journal.pgen.1002796

Liu Q, Wang F, Axtell MJ. Analysis of Complementarity Requirements for Plant MicroRNA Targeting Using a Nicotiana benthamiana Quantitative Transient Assay. Plant Cell. 2014:26(2):741–753. 10.1105/tpc.113.120972

Lu Z, Marand AP, Ricci WA, Ethridge CL, Zhang X, Schmitz RJ. The prevalence, evolution and chromatin signatures of plant regulatory elements. Nat. Plants. 2019:5(12):1250–1259. 10.1038/s41477-019-0548-z

Marand AP, Eveland AL, Kaufmann K, Springer NM. *cis*-Regulatory Elements in Plant Development, Adaptation, and Evolution. Annu. Rev. Plant Biol. 2023:74(1):annurev-arplant-070122-030236. 10.1146/annurev-arplant-070122-030236

Masella AP, Bartram AK, Truszkowski JM, Brown DG, Neufeld JD. PANDAseq: paired-end assembler for illumina sequences. BMC Bioinformatics. 2012:13(1):1–7. 10.1186/1471-2105-13-31

Mcdonald BR, Picard C, Brabb IM, Savenkova MI, Schmitz RJ, Jacobsen SE, Duttke SH. Enhancers associated with unstable RNAs are rare in plants. bioRxiv. 2023. 10.1101/2023.09.25.559415

Nagy F, Boutry M, Hsu MY, Wong M, Chua NH. The 5′-proximal region of the wheat Cab-1 gene contains a 268-bp enhancer-like sequence for phytochrome response. EMBO J. 1987:6(9):2537–2542. 10.1002/j.1460-2075.1987.tb02541.x

O’Malley RC, Huang SC, Song L, Lewsey MG, Bartlett A, Nery JR, Galli M, Gallavotti A, Ecker JR. Cistrome and Epicistrome Features Shape the Regulatory DNA Landscape. Cell. 2016:165(5):1280–1292. 10.1016/j.cell.2016.04.038

Ondek B, Gloss L, Herr W. The SV40 enhancer contains two distinct levels of organization. Nature. 1988:333(6168):40–45. 10.1038/333040a0

Panne D. The enhanceosome. Curr. Opin. Struct. Biol. 2008:18(2):236–242. 10.1016/j.sbi.2007.12.002

Schmitz RJ, Grotewold E, Stam M. Cis-regulatory sequences in plants: Their importance, discovery, and future challenges. Plant Cell. 2022:34(2):718–741. 10.1093/plcell/koab281

Silver BD, Willett CG, Maher KA, Wang D, Deal RB. Differences in transcription initiation directionality underlie distinctions between plants and animals in chromatin modification patterns at genes and cis-regulatory elements. bioRxiv. 2023. 10.1101/2023.11.03.565513

Simpson J, Schell J, Montagu MV, Herrera-Estrella L. Light-inducible and tissue-specific pea lhcp gene expression involves an upstream element combining enhancer- and silencer-like properties. Nature. 1986:323(6088):551–554. 10.1038/323551a0

Song BP, Ragsac MF, Tellez K, Jindal GA, Grudzien JL, Le SH, Farley EK. Diverse logics and grammar encode notochord enhancers. Cell Rep. 2023:42(2):112052. 10.1016/j.celrep.2023.112052

Spitz F, Furlong EEM. Transcription factors: from enhancer binding to developmental control. Nat. Rev. Genet. 2012:13(9):613–626. 10.1038/nrg3207

Thanos D, Maniatis T. Virus induction of human IFNβ gene expression requires the assembly of an enhanceosome. Cell. 1995:83(7):1091–1100. 10.1016/0092-8674(95)90136-1

Thieffry A, Vigh ML, Bornholdt J, Ivanov M, Brodersen P, Sandelin A. Characterization of Arabidopsis thaliana promoter Bidirectionality and Antisense RNAs by Depletion of Nuclear RNA Decay Pathways. Plant Cell. 2020: 10.1105/tpc.19.00815

Tian F, Yang D-C, Meng Y-Q, Jin J, Gao G. PlantRegMap: charting functional regulatory maps in plants. Nucleic Acids Res. 2020:48(D1):D1104–D1113. 10.1093/nar/gkz1020

Tiwari SB, Shen Y, Chang H-C, Hou Y, Harris A, Ma SF, McPartland M, Hymus GJ, Adam L, Marion C, Belachew A, Repetti PP, Reuber TL, Ratcliffe OJ. The flowering time regulator CONSTANS is recruited to the FLOWERING LOCUS T promoter via a unique cis-element. New Phytol. 2010:187(1):57–66. 10.1111/j.1469-8137.2010.03251.x

Uhl JD, Zandvakili A, Gebelein B. A Hox Transcription Factor Collective Binds a Highly Conserved Distal-less cis-Regulatory Module to Generate Robust Transcriptional Outcomes. PLOS Genet. 2016:12(4):e1005981. 10.1371/journal.pgen.1005981

Walcher CL, Nemhauser JL. Bipartite Promoter Element Required for Auxin Response. Plant Physiol. 2012:158(1):273–282. 10.1104/pp.111.187559

Wang X, Aguirre L, Rodríguez-Leal D, Hendelman A, Benoit M, Lippman ZB. Dissecting cis-regulatory control of quantitative trait variation in a plant stem cell circuit. Nat. Plants. 2021:7(4):419–427. 10.1038/s41477-021-00898-x

Weber B, Zicola J, Oka R, Stam M. Plant Enhancers: A Call for Discovery. Trends Plant Sci. 2016:21(11):974–987. 10.1016/j.tplants.2016.07.013

Yan W, Chen D, Schumacher J, Durantini D, Engelhorn J, Chen M, Carles CC, Kaufmann K. Dynamic control of enhancer activity drives stage-specific gene expression during flower morphogenesis. Nat. Commun. 2019:10(1):1705. 10.1038/s41467-019-09513-2

Zenker S, Wulf D, Meierhenrich A, Becker S, Eisenhut M, Stracke R, Weisshaar B, Bräutigam A. Transcription factors operate on a limited vocabulary of binding motifs in Arabidopsis thaliana. bioRxiv. 2023. 10.1101/2023.08.28.555073

