## Supplemental Figures S1 - S15 for "Cooperativity and additivity in plant enhancers"

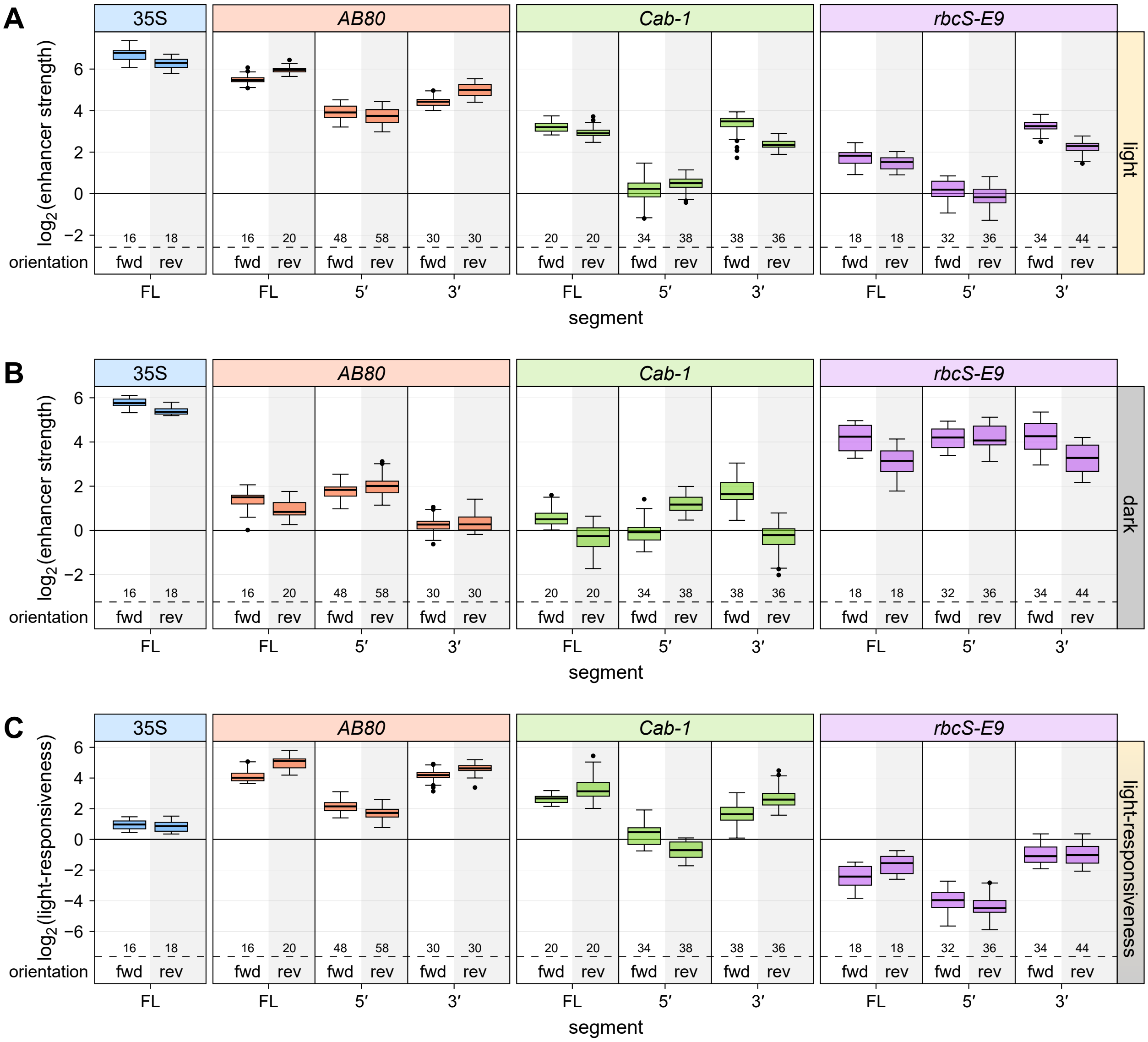


**Supplemental Figure S1** Enhancer strength and light-responsiveness is orientation-independent. (Supports Figure 1) **A** and **B**, Full-length (FL) enhancers, as well as 169-bp long segments from their 5′ or 3′ end, of the *Pisum sativum AB80* and *rbcS-E9* genes and the *Triticum aestivum Cab-1* gene were cloned in the forward (fwd; data reproduced from Figure 1, B and C) or reverse (rev) orientation upstream of the 35S minimal promoter driving the expression of a barcoded GFP reporter gene. All constructs were pooled and the viral 35S enhancer was added as an internal control. The pooled enhancer library was subjected to Plant STARR-seq in tobacco leaves with plants grown for 2 days in normal light/dark cycles (**A**) or completely in the dark (**B**) prior to RNA extraction. Enhancer strength was normalized to a control construct without an enhancer ($\log_2$ set to 0). **C**, Light-responsiveness ($\log_2$[enhancer strength$^{light}$/enhancer strength$^{dark}$]) was determined for the indicated enhancer segments. Box plots represent the median (center line), upper and lower quartiles (box limits), 1.5× interquartile range (whiskers), and outliers (points) for all corresponding barcodes from two replicates. Numbers at the bottom of each box plot indicate the number of barcodes in each group.

Supplemental Data. Jores et al.

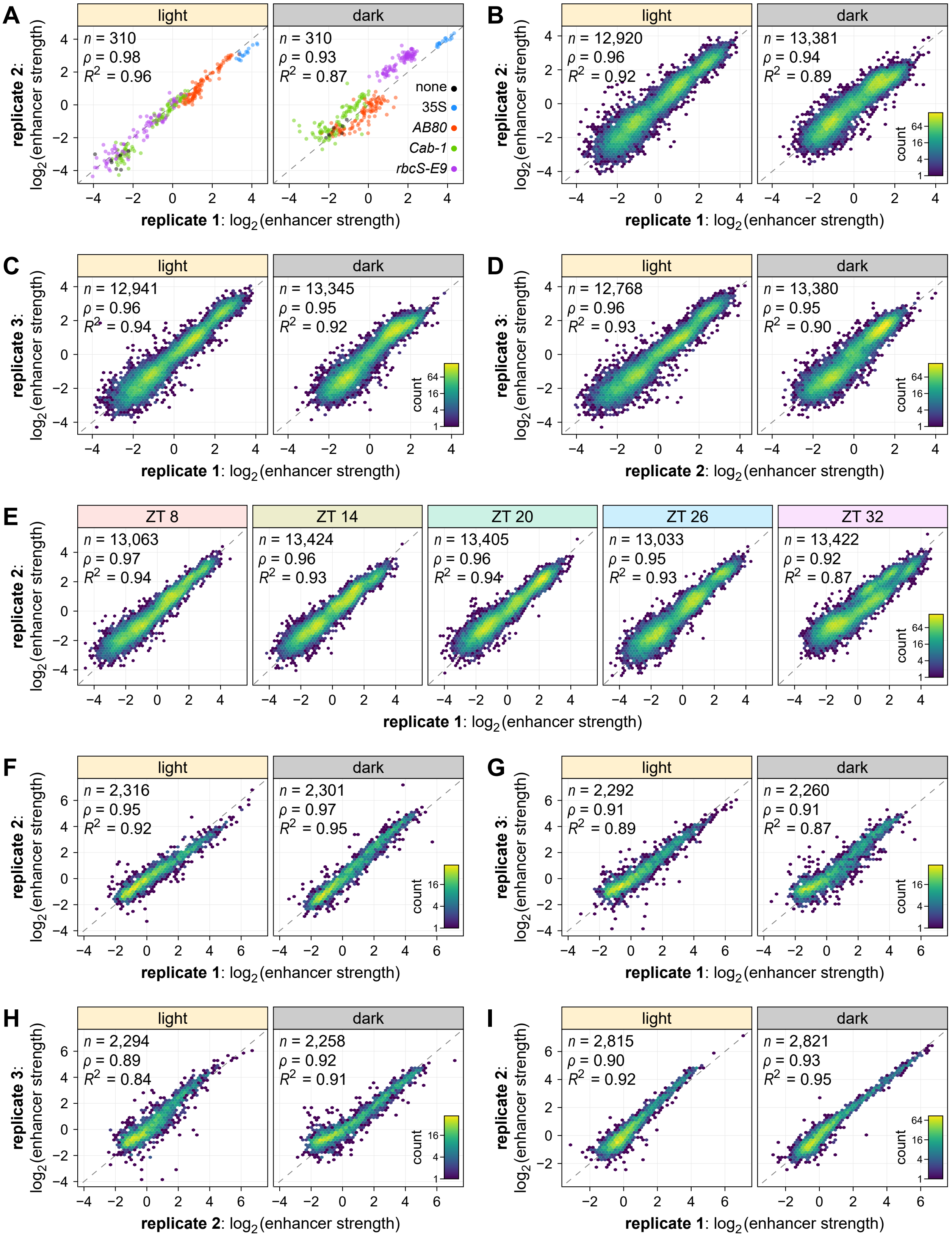

**Supplemental Figure S2** Plant STARR-seq yields highly reproducible results. (Supports all figures) **A**–**I**, Correlation between biological replicates of Plant STARR-seq for the enhancer library used in Figure 1 and Supplemental Figure S1 (**A**), the single-nucleotide enhancer variants library used in Figures 2–4 and Supplemental Figures S4–S13 (**B**–**E**), the synthetic enhancer library used in Figures 6 and 7 and Supplemental Figures S13–S15 (**F**–**H**), and the combined double-deletion and synthetic enhancer valdiation library used in Figure 5 and Supplemental Figure S13 (**I**) performed under the indicated condition or at the indicated time points. Pearson's $R^2$, Spearman's $\rho$, and number ($n$) of enhancer variants or enhancer fragment combinations are indicated. The color in the hexbin plots in (**B**–**I**) represents the count of points in each hexagon.

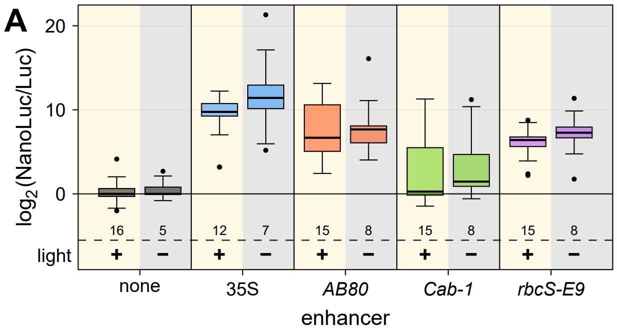


**Supplemental Figure S3** The dual-luciferase assay cannot detect light-responsive enhancer activity. (Supports Figure 1) **A**, Transgenic *Arabidopsis* lines were generated with constructs harboring a constitutively expressed luciferase (Luc) gene and a nanoluciferase (NanoLuc) gene under control of a 35S minimal promoter coupled to the 35S enhancer or the 3′ segments of the *AB80*, *Cab-1*, or *rbcS-E9* enhancers (see Figure 1D). Nanoluciferase activity was measured in 3–5 T2 plants from these lines and normalized to the activity of luciferase. Plants were either grown in normal light/dark cycles (+ light; data reproduced from Figure 1E) or shifted to complete darkness (− light) for 4 days prior to sample collection. The NanoLuc/Luc ratio was normalized to a control construct without an enhancer (none; $\log_2$ set to 0). Box plots represent the median (center line), upper and lower quartiles (box limits), 1.5× interquartile range (whiskers), and outliers (points) for all corresponding plant lines from two (− light) or three (+ light) independent replicates. Numbers at the bottom of each box plot indicate the number of samples in each group.

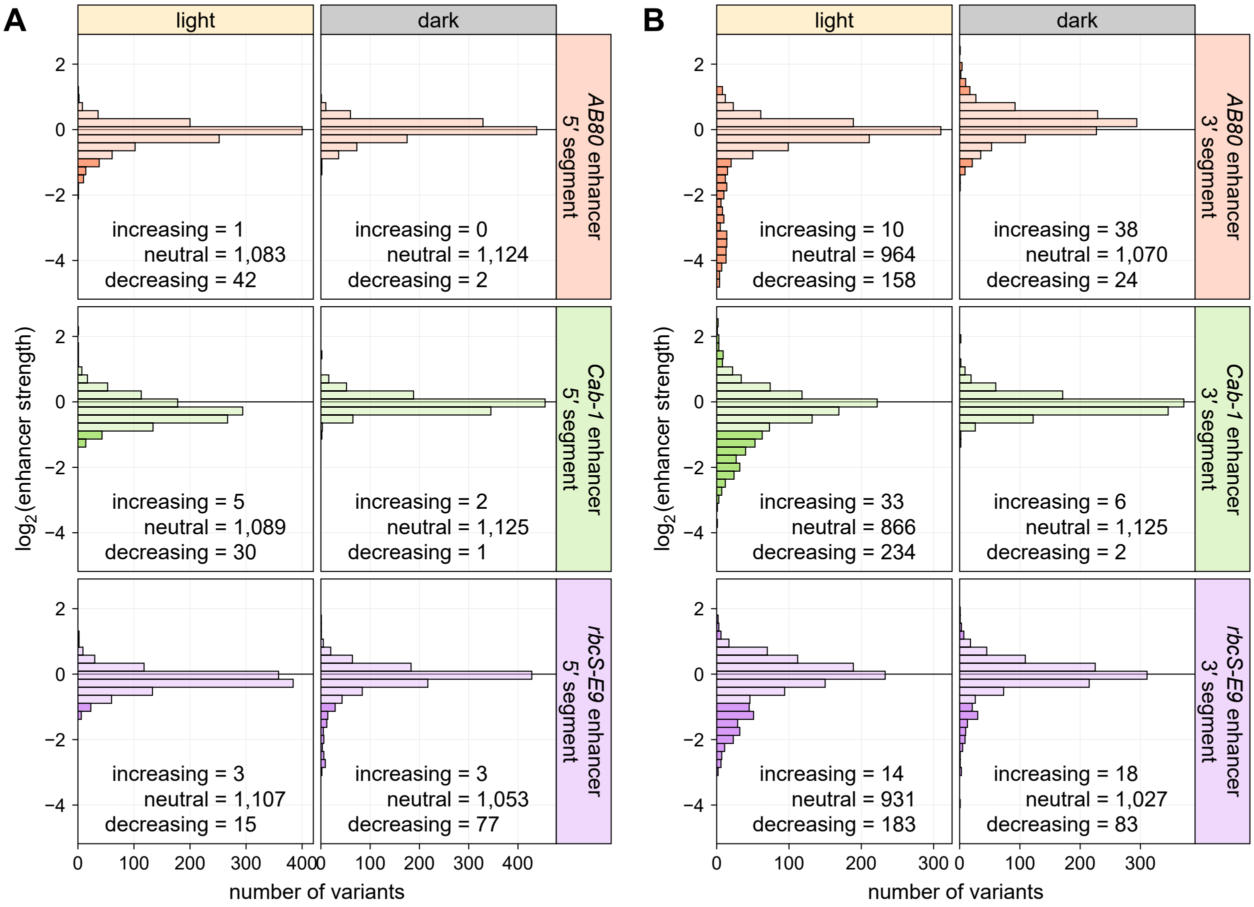


**Supplemental Figure S4** Few single-nucleotide mutations have a strong effect on enhancer strength. (Supports Figures 2 and 3) **A** and **B**, All possible single-nucleotide substitution, deletion, and insertion variants of the 5′ (**A**) and 3′ (**B**) segments of the *AB80*, *Cab-1*, and *rbcS-E9* enhancers were subjected to Plant STARR-seq in tobacco plants grown in normal light/dark cycles (light) or completely in the dark (dark) for two days prior to RNA extraction. Enhancer strength was normalized to the wild-type variant ($\log_2$ set to 0). Variants were grouped into three categories: increasing, $\log_2$(enhancer strength) > 1; neutral, $\log_2$(enhancer strength) between −1 and 1; decreasing, $\log_2$(enhancer strength) < −1. The number of variants in each category is indicated. Neutral variants are shown in a lighter color than increasing and decreasing variants in the histograms.

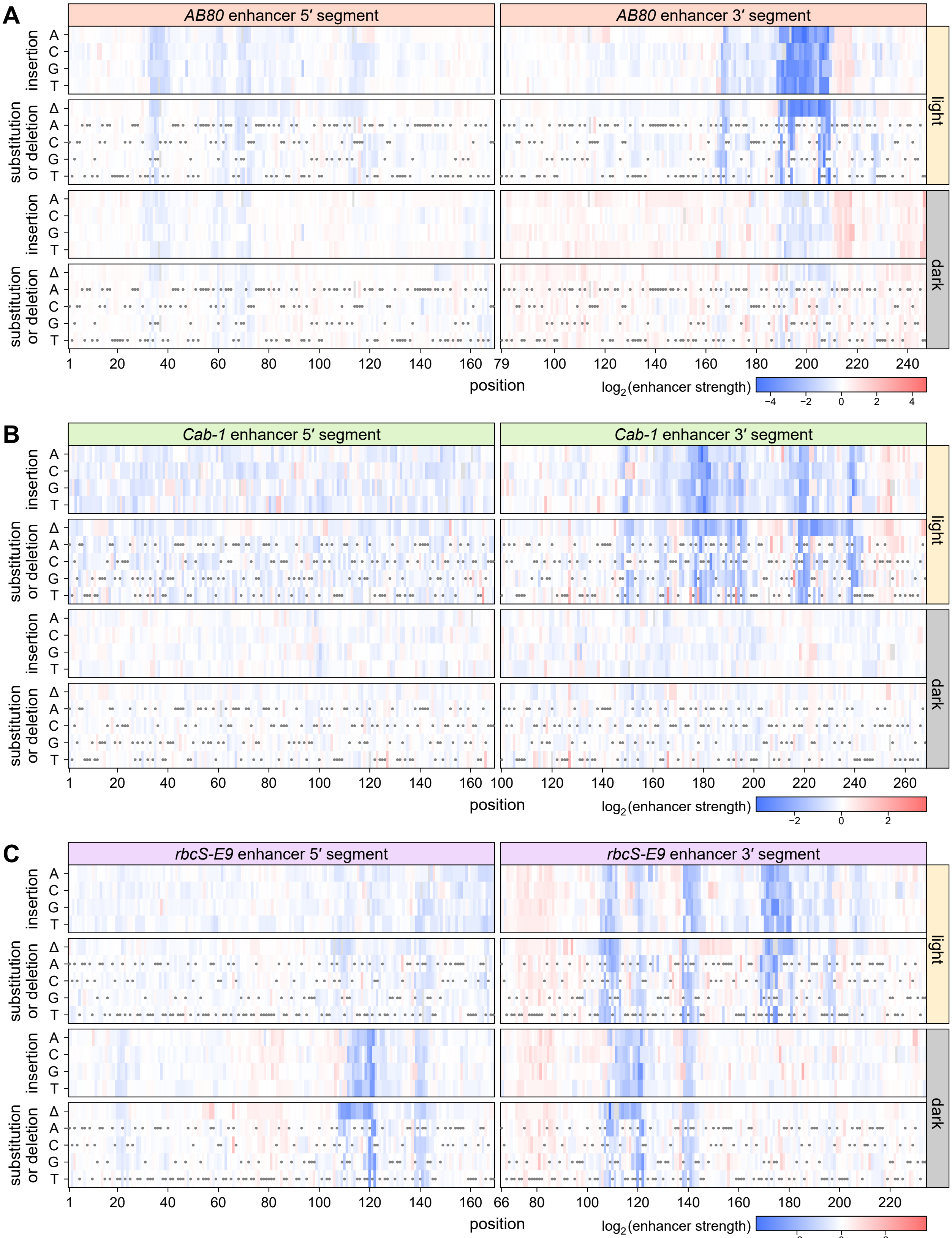
A
AB80 enhancer 5′ segment
AB80 enhancer 3′ segment
insertion
substitution or deletion
A
C
G
T
Δ
light
dark
position
log2(enhancer strength)
−4 −2 0 2 4
B
Cab-1 enhancer 5′ segment
Cab-1 enhancer 3′ segment
−2 0 2
C
rbcS-E9 enhancer 5′ segment
rbcS-E9 enhancer 3′ segment
−2 0 2

**Supplemental Figure S5** Saturation mutagenesis reveals mutation-sensitive patches in plant enhancers. (Supports Figures 2 and 3) **A**–**C**, All possible single-nucleotide substitution, deletion, and insertion variants of the 5′ and 3′ segments of the *AB80* (**A**), *Cab-1* (**B**), and *rbcS-E9* (**C**) enhancers were subjected to Plant STARR-seq in tobacco plants grown in normal light/dark cycles (light) or completely in the dark (dark) for two days prior to RNA extraction. Enhancer strength was normalized to the wild-type variant ($\log_2$ set to 0) and plotted as a heatmap. Missing values are shown in light gray and wild-type variants are marked with a gray dot.

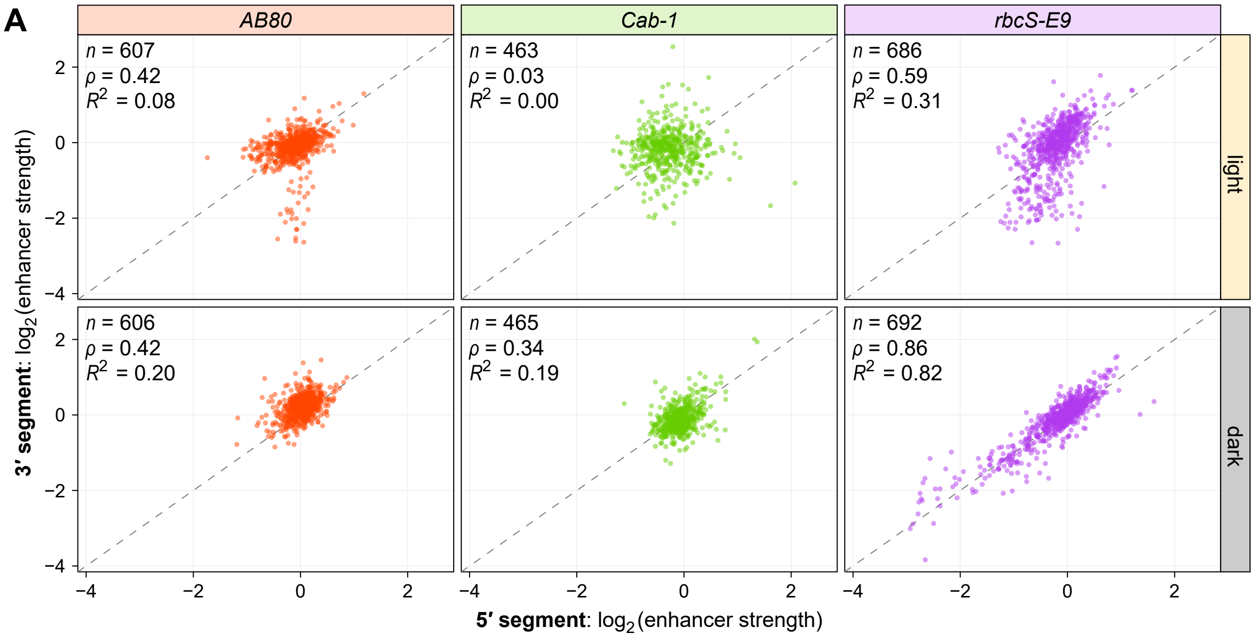


**Supplemental Figure S6** Effects of mutations in the overlap region of the 5′ and 3′ enhancer segments are more similar in the dark. (Supports Figure 2) **A**, All possible single-nucleotide substitution, deletion, and insertion variants of the 5′ and 3′ segments of the *AB80*, *Cab-1*, and *rbcS-E9* enhancers were subjected to Plant STARR-seq in tobacco plants grown in normal light/dark cycles (light) or completely in the dark (dark) for two days prior to RNA extraction. Enhancer strength was normalized to the wild-type variant ($\log_2$ set to 0). For mutations located in the overlap region between the two segments (positions 79–169 in *AB80*, positions 100–169 in *Cab-1*, and positions 66–169 in *rbcS-E9*), the normalized enhancer strength measured in the context of the 5′ segment (x axis) is compared against the normalized enhancer strength measured in the context of the 3′ segment (y axis). Pearson's $R^2$, Spearman's $\rho$, and number ($n$) of enhancer variants are indicated.

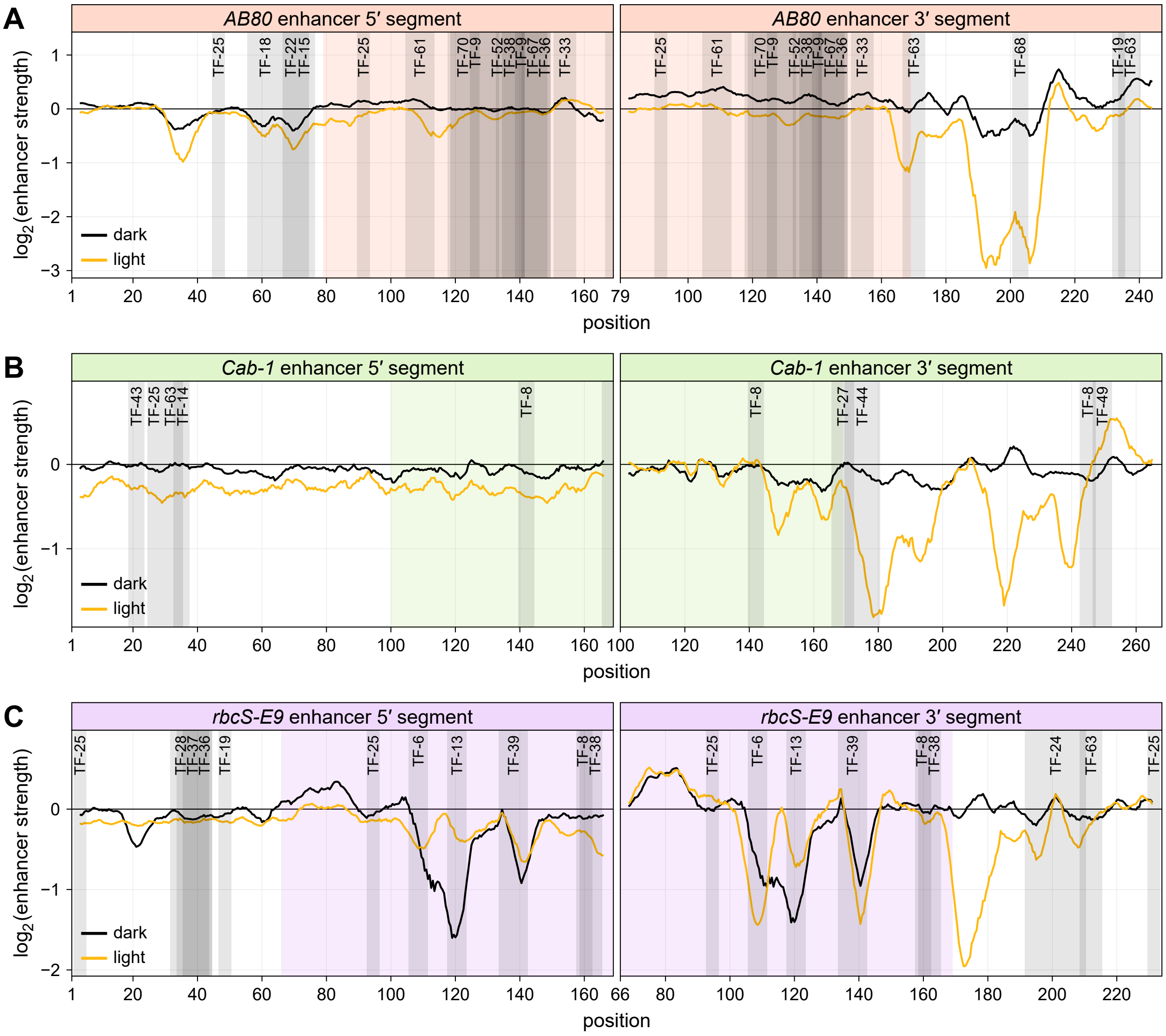


**D**

| ID | family | consensus | ID | family | consensus |
|---|---|---|---|---|---|
| TF-6 | bHLH/BES1/bZIP/Trihelix | CACGTG | TF-36 | HD-ZIP/YABBY | ATAATAATw |
| TF-8 | Dof/C3H | AAAAG | TF-37 | WOX/HD-ZIP | CAATCATAws |
| TF-9 | ZF-HD/HD-ZIP | ATATTdrnnnnnwwwT | TF-38 | NF-YB/C2H2/G2-like | TTGAAAA |
| TF-13 | MYB related/MYB/GeBP | GGATAA | TF-39 | HD-ZIP/YABBY | TAATCATTA |
| TF-14 | SBP/AP2/C2H2/bHLH | CGTAC | TF-43 | E2F/DP | GCGCC |
| TF-15 | TCP | TGGGCCCACn | TF-44 | Trihelix | TAACCATGTTr |
| TF-18 | C2H2/GRAS | AAAGACAAAAm | TF-49 | M-type MADS | TCACCA |
| TF-19 | GATA | GATC | TF-52 | B3 | AGAAAnwnnnAAGAAAn |
| TF-22 | TCP | GGGACCAC | TF-61 | C2H2 | rAAACAGAG |
| TF-24 | GATA/MIKC MADS | CATCATCATCATCATCATC | TF-63 | NF-YB | TCCATCA |
| TF-25 | C2H2 | CACT | TF-67 | B3 | AAAAAAAAAAA |
| TF-27 | MYB related/Trihelix | TTAGGGy | TF-68 | Nin-like | CAGCA |
| TF-28 | CPP | TTTAATTTTrAww | TF-70 | WOX | AnTTAATTA |
| TF-33 | RAV | TmTGTTG | | | |

**Supplemental Figure S7** Mutation-sensitive regions contain few strong matches to known transcription factor binding motifs. (Supports Figure 3) **A**–**C**, The wild-type sequences of the 5′ and 3′ segments of the *AB80* (**A**), *Cab-1* (**B**), and *rbcS-E9* (**C**) enhancers were scanned for significant matches to known transcription factor binding motifs. Hits are shown as gray areas overlaid on the mutational sensitivity plots reproduced from Figure 2. The ID of the matching transcription factor motif is indicated. **D**, For each transcription factor motif ID, the table lists the families of transcription factors that can bind to it and the consensus binding site sequence. Ambiguous nucleotides in the consensus sequence are shown as lowercase and correspond to: n = A, C, G, or T; d = A, G, or T; r = A or G; w = A or T; m = A or C.

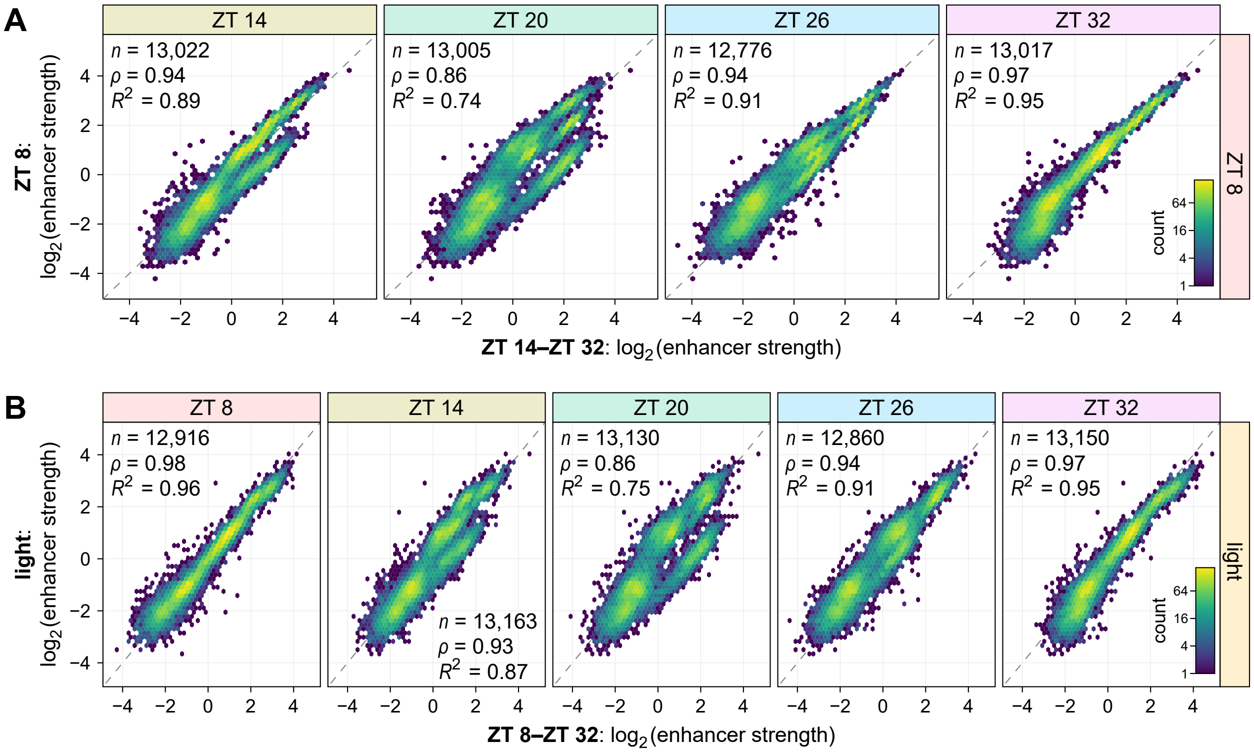


**Supplemental Figure S8** Strong correlation between Plant STARR-seq samples obtained 24 hours apart from each other. (Supports Figure 4) **A**, All possible single-nucleotide variants of the *AB80*, *Cab-1*, and *rbcS-E9* enhancers were subjected to Plant STARR-seq in tobacco leaves. On the morning of the third day after transformation (ZT 0), the plants were shifted to constant light. Leaves were harvested for RNA extraction starting at mid-day (ZT 8) and in 6 hour intervals (ZT 14, 20, 26, and 32) afterwards. Hexbin plots (color represents the count of points in each hexagon) of the correlation between samples obtained at the indicated time points are shown. Pearson's $R^2$, Spearman's $\rho$, and number ($n$) of enhancer variants are indicated. **B**, Hexbin plots of the correlation between the samples from the time course experiment described above compared to the same library tested in normal light/dark cycles (light) as described in Figure 2. The "light" samples were harvested at ZT 8.

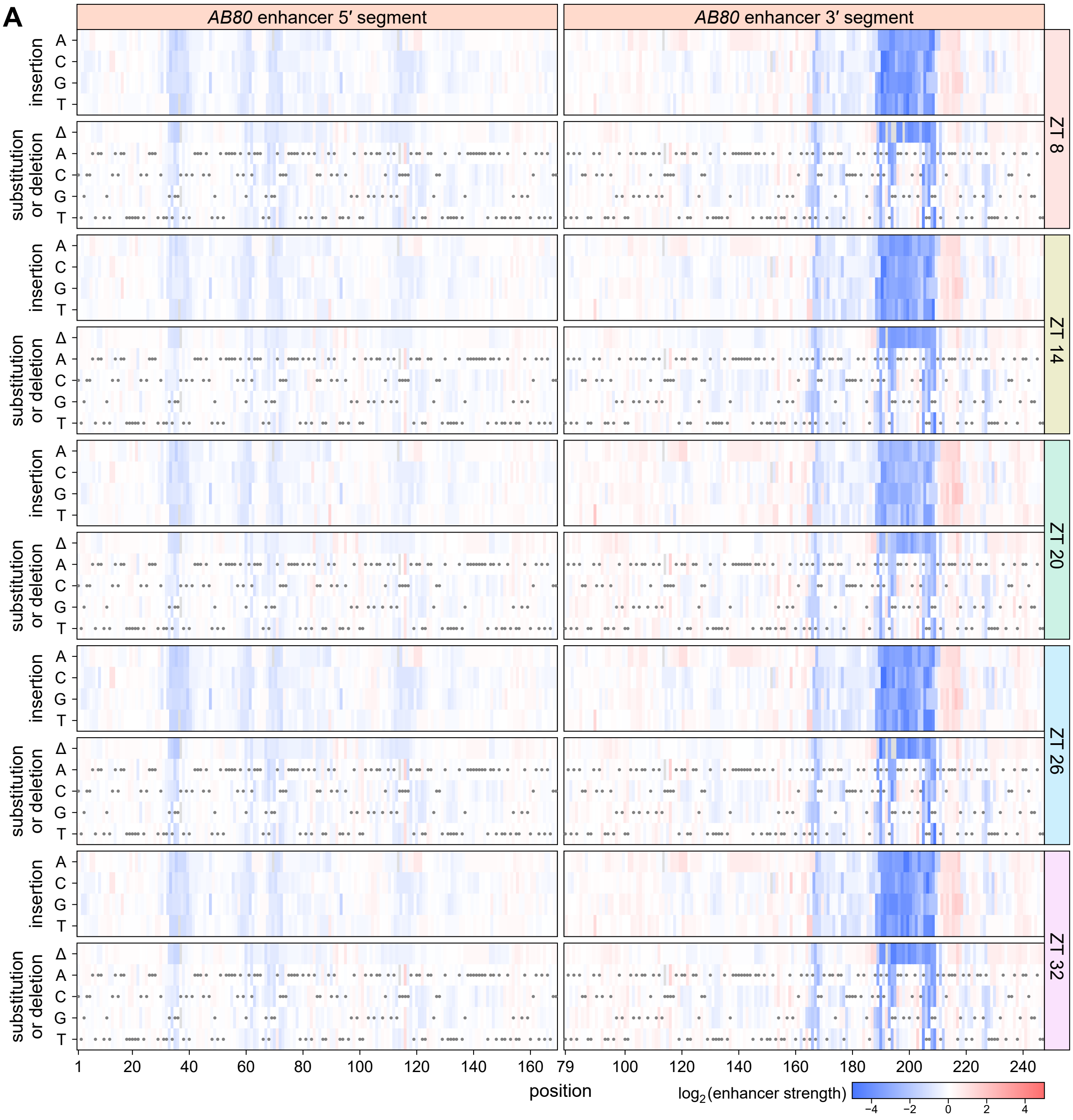


**Supplemental Figure S9** Saturation mutagenesis maps of the *AB80* enhancer do not change much over time in constant light. (Supports Figure 4) **A**, All possible single-nucleotide variants of the *AB80* enhancer were subjected to Plant STARR-seq in tobacco leaves. On the morning of the third day after transformation (ZT 0), the plants were shifted to constant light. Leaves were harvested for RNA extraction starting at mid-day (ZT 8) and in 6 hour intervals (ZT 14, 20, 26, and 32) afterwards. Enhancer strength was normalized to the wild-type variant ($\log_2$ set to 0) and plotted as a heatmap. Missing values are shown in light gray and wild-type variants are marked with a gray dot.

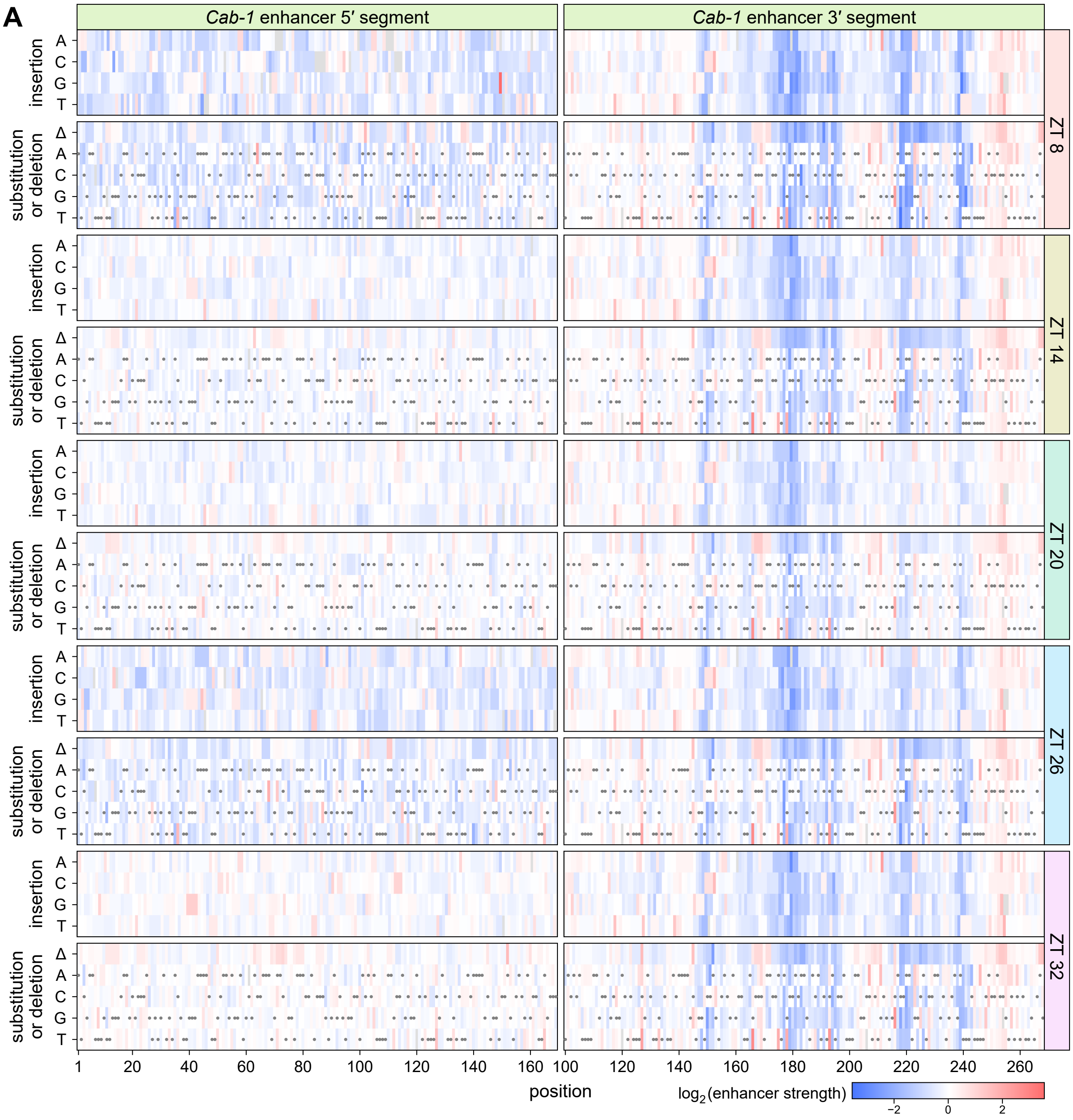


**Supplemental Figure S10** Saturation mutagenesis maps of the *Cab-1* enhancer do not change much over time in constant light. (Supports Figure 4) **A**, The *Cab-1* enhancer was subjected to the same experiment as the *AB80* enhancer in Supplemental Figure S9.

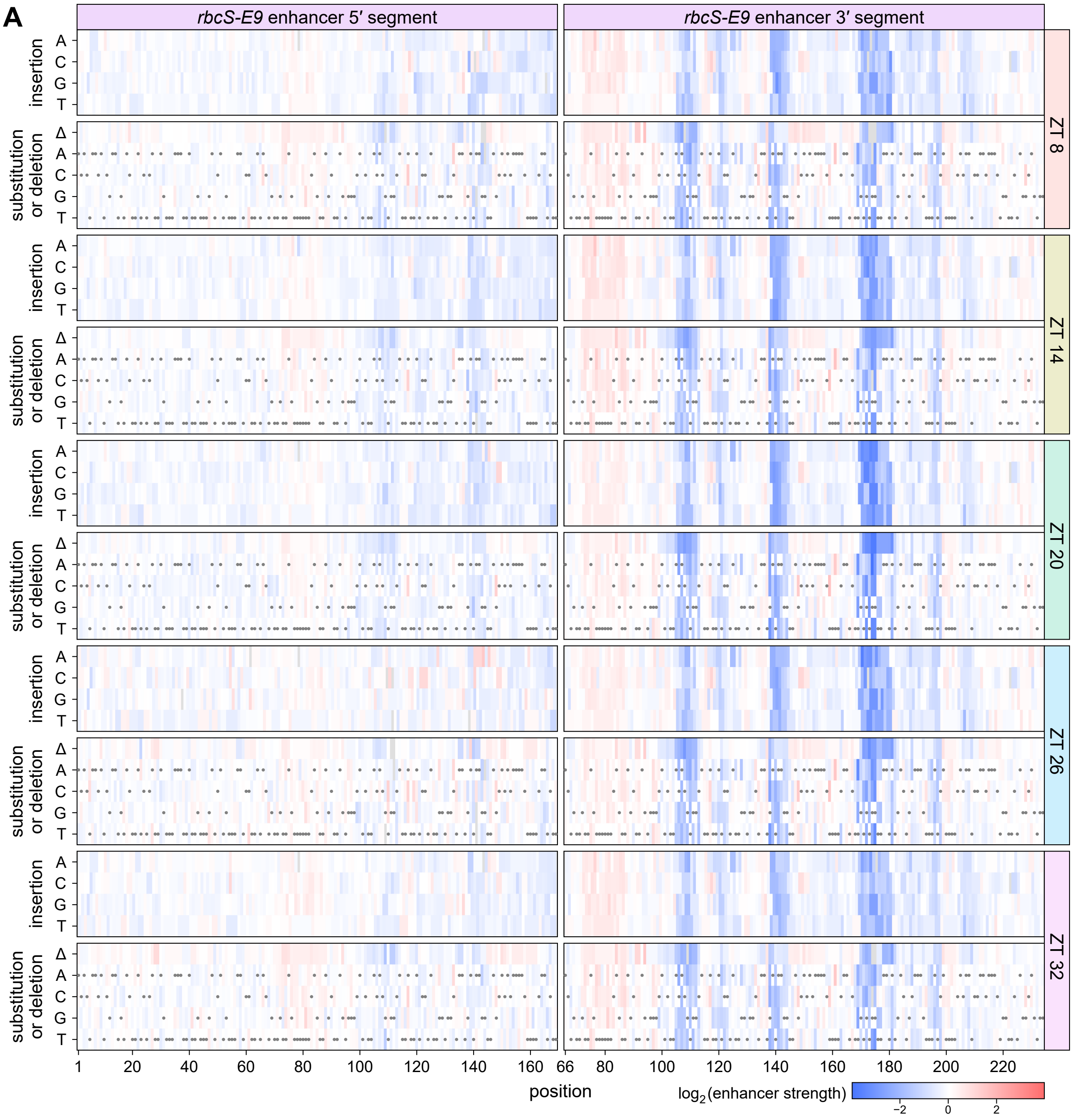


**Supplemental Figure S11** Saturation mutagenesis maps of the *rbcS-E9* enhancer do not change much over time in constant light. (Supports Figure 4) **A**, The *rbcS-E9* enhancer was subjected to the same experiment as the *AB80* enhancer in Supplemental Figure S9.

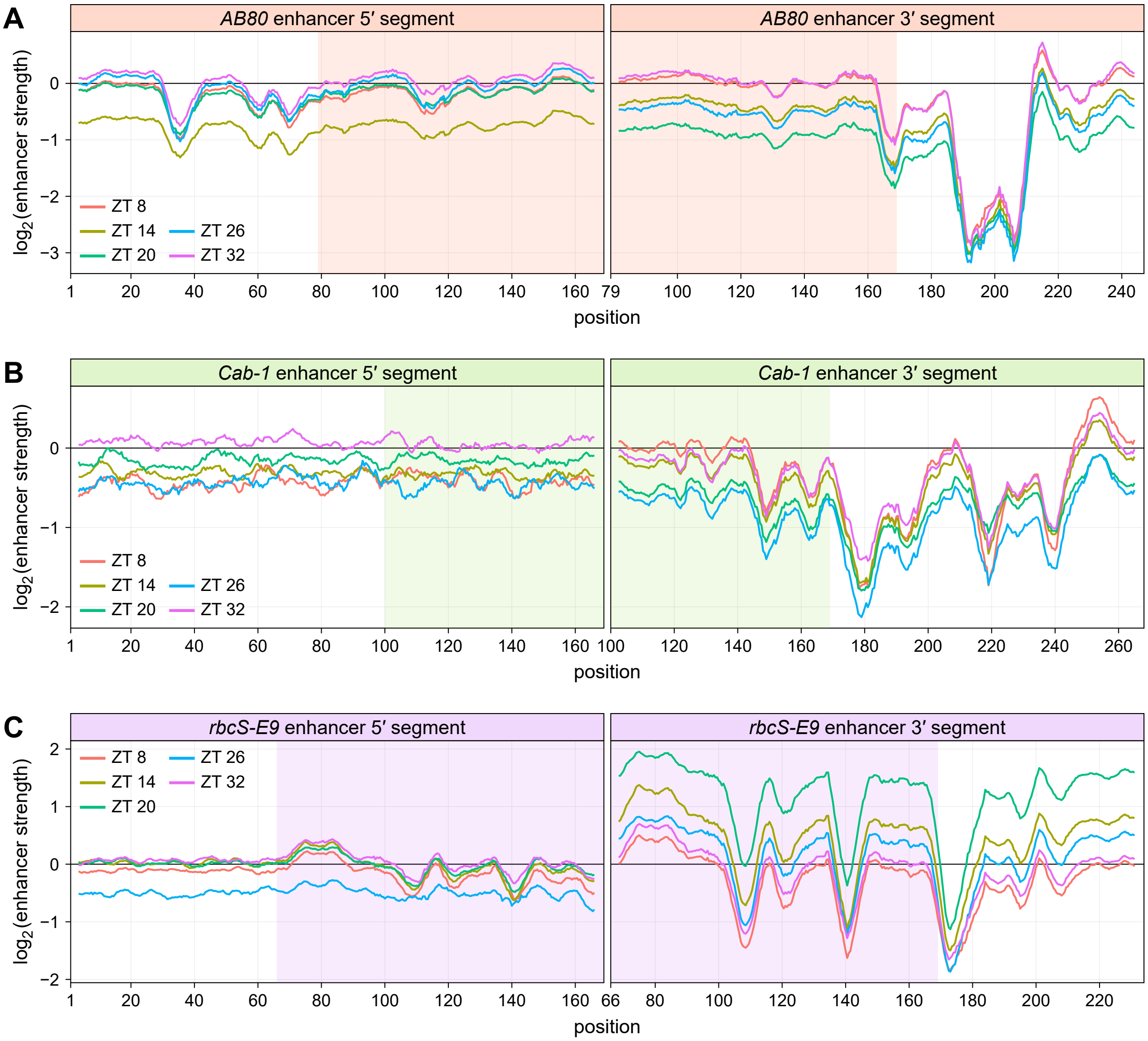


**Supplemental Figure S12** Mutation-sensitive regions of the *AB80*, *Cab-1*, and *rbcS-E9* enhancers are conserved over time in constant light. (Supports Figure 4) **A**–**C**, All possible single-nucleotide substitution, deletion, and insertion variants of the 5′ and 3′ segments of the *AB80* (**A**), *Cab-1* (**B**), and *rbcS-E9* (**C**) enhancers were subjected to Plant STARR-seq in tobacco leaves. On the morning of the third day after transformation (ZT 0), the plants were shifted to constant light. Leaves were harvested for RNA extraction starting at mid-day (ZT 8) and in 6 hour intervals (ZT 14, 20, 26, and 32) afterwards. Enhancer strength was normalized to the wild-type variant ($\log_2$ set to 0). A sliding average (window size = 6 bp) of the mean enhancer strength for all variants at a given position is shown. The shaded area indicates the region where the 5′ and 3′ segments overlap.

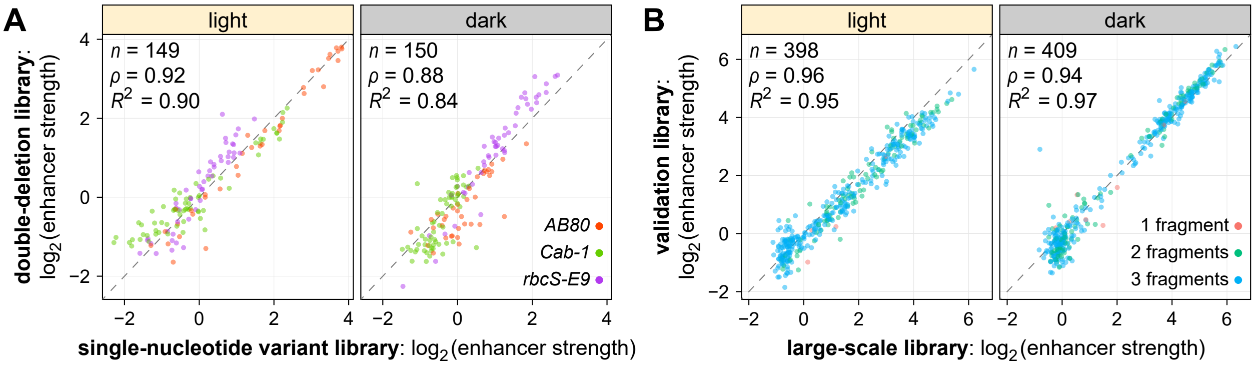


**Supplemental Figure S13** Plant STARR-seq experiments are reproducible across libraries. (Supports Figures 2, 3 and 5–7) **A**, Correlation between the enhancer strength of single-nucleotide deletion variants of the *AB80*, *Cab-1*, and *rbcS-E9* enhancers present in the comprehensive single-nucleotide enhancer variants library (described in Figure 2) and in a second, independent library with single- and double-deletion enhancer variants (described in Figure 5). **B**, Correlation between the strength of synthetic enhancers created by combining fragments of the *AB80*, *Cab-1*, and *rbcS-E9* enhancers as measured in the large-scale library (described in Figure 6) and in a second, smaller validation library. Pearson's $R^2$, Spearman's $\rho$, and number ($n$) of enhancer variants are indicated.

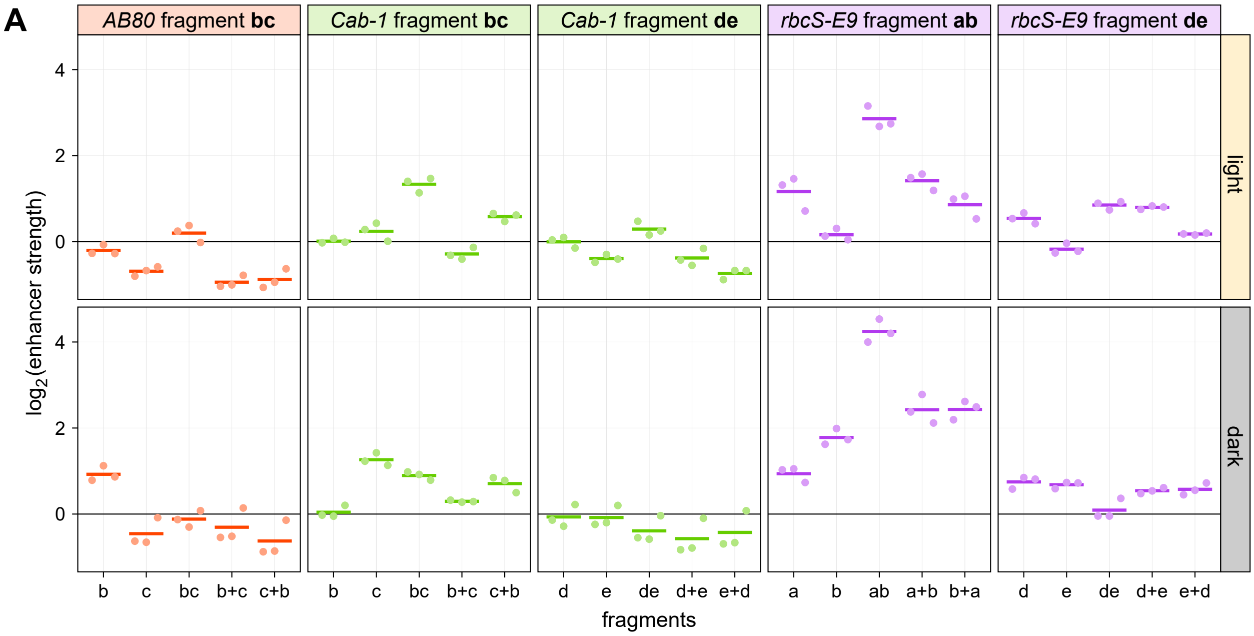


**Supplemental Figure S14** Correct spacing between mutation-sensitive regions is required for full activity. (Supports Figure 6) **A**, Plots of the strength in the indicated condition of enhancer fragments (Figure 6D) or fragment combinations (separated by a + sign and shown in the order in which they appear in the construct; Figure 6E) in three replicates (points) and the mean strength (lines). Enhancer strength was normalized to a control construct without an enhancer ($log_2$ set to 0). Some plots are reproduced from Figure 6, F and G.

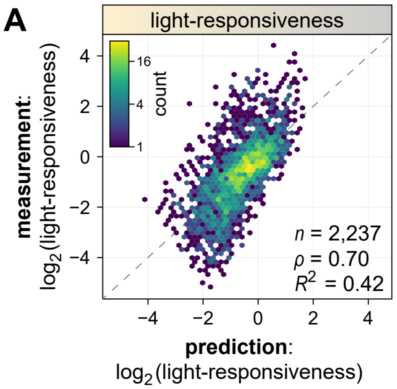


**Supplemental Figure S15** A linear model can predict the light-responsiveness of synthetic enhancers. (Supports Figure 7) **A**, A linear model was built to predict the light-responsiveness of synthetic enhancers created by randomly combining up to three fragments derived from mutation-sensitive regions of the *AB80*, *Cab-1*, and *rbcS-E9* enhancers (see Figure 6A) based on the light-responsiveness of the constituent individual fragments. A hexbin plot (color represents the count of points in each hexagon) of the correlation between the model's prediction and the measured data is shown. Pearson's $R^2$, Spearman's $\rho$, and number ($n$) of enhancer fragment combinations are indicated.
